# Altered global modular organization of intrinsic functional connectivity in autism arises from atypical node-level processing

**DOI:** 10.1101/2022.07.30.502167

**Authors:** Priyanka Sigar, Lucina Q. Uddin, Dipanjan Roy

## Abstract

Autistic spectrum disorder (ASD) is a neurodevelopmental condition characterized by restricted interests and repetitive behaviors as well as social-communication deficits. These traits are associated with atypicality of functional brain networks. Modular organization in the brain plays a crucial role in network stability and adaptability for neurodevelopment. Previous neuroimaging research demonstrates discrepancies in studies of functional brain modular organization in ASD. These discrepancies result from the examination of mixed age groups. Furthermore, recent findings suggest while much attention has been given to deriving atlases and measuring the connections between nodes, the within nodes information may be crucial in determining altered modular organization in ASD compared with TD. However, altered modular organization originating from systematic nodal changes are yet to be explored in younger children with ASD. Here, we used graph-theoretical measures to fill this knowledge gap. To this end, we utilized multicenter resting-state BOLD fMRI data collected from 5–10-year-old children - 34 ASD and 40 typically developing obtained from the Autism Brain Image Data Exchange (ABIDE) I and II. We demonstrated alterations in the topological roles and modular cohesiveness are the two key properties of the brain regions anchored in default mode, sensorimotor, and salience networks primarily relates to social and sensory deficits in ASD children. These results demonstrate atypical global network organization in ASD children arise from nodal role changes and contribute to the growing body of literature suggesting that there is interesting information within nodes providing critical marker of functional brain networks in Autistic children.

## INTRODUCTION

Graph-theoretical approaches applied to magnetic resonance imaging (MRI) data have revealed that the human brain exhibits a hierarchical modular organization, with relatively larger functional communities further divisible into smaller communities. (Sporns and Betzel 2016; Meunier, Lambiotte, and Bullmore 2010). Within these modular partitions, each brain region (node) has its distinct functional role in information processing within and across different modules, determined by a specific profile of within- and between module connectivity. This modular profile of nodes helps classify them into different node types, whose relative properties may affect information flow within a complex network system (Sporns 2014). Identification of functional brain modular structures can be used to delineate functional components associated with specific biological functions (Hsu et al., 2012). Modular structures play a crucial role in network stability and adaptability to facilitate optimal network functioning (He et al., 2009, Bassett & Gazzaniga, 2011). The modular organization of the brain may play a crucial role in evolution and neurodevelopment (D et al., 2009).

Autism spectrum disorder (ASD) is a neurodevelopmental condition characterized by restricted interests and repetitive behaviors as well as difficulty with social-communication skills. Recent static functional connectivity (SFC) analysis of resting-state functional magnetic resonance imaging (rsfMRI) data has shown atypical functional connectivity and functional brain network configurations associated with social cognitive abilities in individuals with ASD (Rudie et al. 2012, 2013). Previous resting-state and task fMRI studies have also reported links between various cognitive functions and modular organization of functional brain networks. Associations have been observed between cognitive abilities (e.g., intelligence, working memory performance, social and emotional processing) and (a) modular network organization (Cohen et al. 2016; Hilger 2020; Hilger et al. 2017; Stevens et al. 2012), (b) proportions of specific node types (i.e., connector hubs, provincial hubs) and (c) alteration in topological roles of brain regions (Cohen et al. 2016 Anon n.d.; Hilger 2020; Hilger et al. 2017; Stanley et al. 2014). Recently, Glerean and colleagues (Glerean et al., 2015) reported group differences (age 19-47 years) in the composition of the default mode network (DMN) and a ventral-temporal-limbic (VTL) sub-network (amygdala, striatum, thalamus, parahippocampal gyrus (PhG), fusiform gyrus (FuG), and inferior frontal gyrus). These findings were significantly correlated with autism symptom severity scores. These studies strongly suggest that individual differences in the modular organization and the topological roles of the nodes of functional brain networks are related to behavioral traits in ASD.

Previous studies explored various large-scale and region-specific modular properties, which included subjects with a wide range of ages, stratifying individuals into different age cohorts (e.g., mixed children and adolescents (4-17 years)), resulting in mixed findings throughout the autism neuroimaging literature (Nomi & Uddin, 2015; Uddin, Supekar, & Menon, 2013). Between the age range of 5-10, hubs shift from being localized in motor areas and primary sensory areas to a more distributed pattern across frontal, temporal, visual, and subcortical regions (Supekar, Musen, and Menon 2009). This shift of hubs reflects the development of higher cognitive order during this period (Oldham et al., 2019). However, atypicality in the large-scale and regional modular properties in younger children with ASD remains unexplored. Here, we address fundamental questions about the influence of alteration in topological roles of brain regions on global modular properties of the brain in younger children with ASD (5-10 years). An outstanding question is (a) whether, in children with ASD, the alteration of global modular structure arises from the atypical modular cohesiveness of specific brain regions (nodes). (b) The alteration in the topological roles of specific brain regions is responsible for atypical modular configuration and functional connectivity in ASD.

In this study, we applied a graph-theoretical approach to resting-state fMRI data from a sample of high-functioning children with ASD and typically developing (TD) children available through the Autism Brain Imaging Exchange (ABIDE) (A. Di Martino et al. 2014; Adriana Di Martino et al. 2017). We included children between 5-10 years of age who were male, right-handed, had full-scale IQ > 75 from multiple datasets, including NYU, Stanford, and San Diego, to test the hypothesis that there would be significant between-group differences in the modular configuration of sensory-motor and neurocognitive networks between ASD and TD children, and that alterations in nodal roles would be linked to autism symptom severity.

## METHODS

### 1. Participants

We used resting-state fMRI data collected from children with ASD (5-10 years; males; right-handed; IQ > 75) from all ABIDE I & II sites (NYU, Stanford, San Diego) (A. Di Martino et al. 2014; Adriana Di Martino et al. 2017) (http://fcon_1000.projects.nitrc.org/indi/abide/) that had participants in the age range 5-10. To create a well-matched TD comparison dataset, we tested for significant group differences in age, full-scale IQ, and framewise displacement using t-tests. An ANOVA was used to test site and group interactions with age, FIQ, and framewise displacement (FD) (**Table.1**).

**Table 1:** Participants demographics.

|  | ASD (n=57) | TD (n=62) | Group comparisons (P-value) |
| --- | --- | --- | --- |
| Age | 9.36 ( ± 1.74) | 9.67( ± 1.54) | 0.31a |
| Site x group interaction |  | - | 0.25b |
| Full-scale IQ | 110.43 ( ± 16.83) | 114.46 ( ± 13.42) | 0.21a |
| Site x group interaction |  | - | 0.74b |
| Mean FD (mm) | 0.14( ± 0.04) | 0.12( ± 0.04) | 0.11a |
| Site x group interaction |  |  | 0.32b |
| ADOS score |  |  |  |
| Total score | 12.15 ( ± 4.21) | - |  |
| Social | 9.36 ( ± 3.44) | - |  |
| RRB score | 2.78 ( ± 1.70) | - |  |
| Severity score | 7.01( ± 1.77) | - |  |

ADOS, Autism Diagnostic Observation Schedule; RRB, Restricted and Repetitive Behaviors. FD, Frame wise displacement. a two-sample t-test; b ANOVA

To further eliminate confounds, we excluded subjects based on the following criteria: (1) high levels of head motion (maximum motion >2mm or 2°rotation, or more than 50% of frames with high FD; time points whose FD was larger than 0.5 mm, along with the preceding time-point and following two time-points, were defined as high head motion time points); (2) incomplete cortical coverage in the scan; (3) age above 3 SD+/- mean across the samples; (4) FIQ above 2SD+/- mean across the samples; (5) sites with less than 10 subjects.

### 2. Data selection and preprocessing

All resting-state fMRI data were preprocessed using the Data Processing Assistant for Resting-State fMRI (DPARSF) toolbox (http://rfmri.org/DPARSF) (Chao-Gan and Yu-Feng, 2010). The initial ten volumes were dropped to ensure steady-state longitudinal magnetization. The functional images were realigned using a six-parameter (rigid body) linear transformation. Structural images (T1- weighted MPRAGE) were segmented into grey matter (GM), white matter (WM), and cerebrospinal fluid (CSF) and co-registered to the mean functional images using a 6 degree of freedom linear transformation. Subsequently, structural images were transformed into standard Montreal Neurological Institute (MNI) space at the resolution of 3 mm^3^ isotropic voxels using the Diffeomorphic Anatomical Registration Through Exponentiated Lie algebra (DARTEL) tool (Ashburner, 2007). The following nuisance signals were removed: head motion effects (Friston 24-parameters) (Friston et al., 1995), signals from WM and CSF, and linear and quadratic trends. WM and CSF signals were regressed using an anatomical component-based noise correction procedure (aCompCor) (Behzadi, Restom, Liau, & Liu, 2007). We did not use global signal regression (GSR) since it has been shown to influence anti-correlations in resting-state brain networks and distort group differences in intrinsic functional connectivity (Murphy, Birn, Handwerker, Jones, & Bandettini, 2009; Weissenbacher et al., 2009). Finally, a temporal bandpass filter (0.01–0.1Hz) was applied to the BOLD time series h (X. Liu & Duyn, 2013). All normalized images were smoothed with a Gaussian kernel to 6 mm full-width at half-maximum (FWHM). Given that head motion-based artifacts influence BOLD signals (Power et al., 2014), we performed covariate regressions of Friston 24-motion parameters using the DPARSF toolbox. To further minimize the impact of head motion artifacts, the scrubbing method was applied, wherein, frames/scans with framewise displacement (FD) >0.5 mm were discarded (flagged frame as well as one before and two after) (Power et al., 2012).

### 3. Construction of resting-state functional connectivity matrix

Several studies suggest that graph properties like modularity are strongly influenced by different graph densities (Ginestet, Nichols, Bullmore, & Simmons, 2011). We thresholded our correlation matrix over a range of thresholds, retaining the strongest 15, 20, 25, 30, 35, 40, and 45% edges. This resulted in seven thresholded graphs for each subject (refer to Supplementary methods). To parcellate the brain into various regions of interest (ROIs), we used the Schaefer-Yeo 2018 atlas with 200 parcellations (Schaefer et al., 2018) (refer to Supp. methods to parcellations 400 & 600). We used the DPARSF toolbox to extract the BOLD time series of each brain region and to construct a subject-specific N x N connectivity matrix, which contains the functional connectivity between each pair of nodes/brain regions. Functional connectivity between brain regions was estimated by calculating Pearson’s correlation coefficient values of the BOLD signal between the BOLD time series. These values were then normalized to z values using Fisher’s z transformation.

### 4. Graph-theoretical analysis

#### 4.1 Modular organization measures

To choose the adequate community detection algorithm for our study and to assess the reliability of our findings across different methods, we used three community detection algorithms - Louvain (Blondel, Guillaume, Lambiotte, & Lefebvre, 2008), Newman- Girvan (Newman & Girvan, 2004), and Consensus agreement (Lancichinetti & Fortunato, 2012). These algorithms were applied separately on seven thresholded graphs and the resulting graph metrics were averaged for each participant.

We explored three whole-brain measures of the functional network modular organization for each subject: global modularity Q, number of modules, and module size. The Louvain method and consensus agreement method showed similar results; however, the Louvain method provided consistent results across all the participants for all three modularity measures. Thus, the Louvain method was used for further analyses and performed 100 optimization runs on each subject. Statistical analysis on subject-specific values (obtained by averaging across results of all the threshold-defined graphs) for these whole-brain modularity measures was conducted using permutation testing (10,000 iterations, p < 0.05, FDR corrected).

#### 4.2 Modular composition analyses

##### 4.2.1) Group-level modularity composition analyses

The modularity measures can only reflect differences in network modularity; these measures cannot reveal differences in modular composition between groups. Normalized mutual information (NMI) metrics (Kuncheva & Hadjitodorov, 2004) were used to compute group-level similarity in the whole-brain community structure between the ASD and TD groups. The NMI function in the Brain Connectivity toolbox was used (Rubinov & Sporns, 2010). This tool computes an NMI-based similarity value, ranging from zero to one, wherein a similarity value closer to one represents identical community affiliation/assignment (refer to Supplementary detailed methods).

##### 4.2.2) Subject-level modularity composition analyses

The above investigation does not allow statistical qualification of modular structure variability within and between groups. When partitions are compared across two subjects, even if two modules might appear quite similar, they may not have the same labels. In other words, a community detection algorithm might assign different labels to different modules across subjects. This problem can be solved either by manual intervention or by overlapping modules with the same label while preserving the differences in modular partitions between groups. Using the NMI approach, we can study the difference in community structure within and between groups at the individual subject-level and node-level (refer to Supplementary methods).

### 4.3) Within- and Between- Module connectivity metrics

An altered community structure can be indicative of changes in the integration and segregation of functional networks, which can be explored by uncovering the intra- and inter-module connectivity patterns of individual nodes. The two measures proposed by (Guimerà & Amaral, 2005) -participation coefficient *(pc)* and within-module degree *(z)* were used to characterize the connectivity of nodes within and between modules. These two-graph metrics were calculated for binarized and proportionally thresholded graphs using seven different cut-offs (15%, 20%, 25%, 30%, 35%, 40% or 45% strongest edges). For each participant, graph metrics were averaged across these six thresholds to generate individual mean maps. For visualization purposes, individual mean *pc***-** and ***z*** - values of each node were averaged across participants. A two-sample t-test was used to compare these graph metrics between the ASD and TD groups, followed by FDR correction.

### 4.4) Node-type cartographic categorization

(Guimerà & Amaral, 2005) classified nodes into seven hubs and non-hubs based on their within- and between module connectivity profiles. Hubs were classified as Provincial (strong within module connectivity, weak between module connectivity), Connector (strong between-network, weak within-network connectivity), and Kinless (equal within- and between- module connectivity). In contrast, non-hubs were classified as ultra-peripheral (strongest within module connectivity), Peripheral (contains most connections within the same module), Nonhub connectors (many links with nodes in other modules), Nonhub kinless (most links with nodes in other modules). We classified nodes as hubs and non-hubs based on within-module degree values, z ≥ 1 as hubs, and z < 1 as non-hubs. Non-hubs were further classified into subtypes based on participation coefficient values, ultra-peripheral (*pc* ≤ 0.05), peripheral (0.05≤ *pc* ≤ 0.62), non-hub connector (0.62 < *pc* ≤ 0.80), and non-hub kinless nodes (pc > 0.80), whereas hubs were divided into three subtypes - provincial (pc ≤ 0.30), connector (0.30 < pc ≤ 0.75), or kinless hubs (pc > 0.75). Proportions of node types were also calculated for binarized and proportionally thresholded graphs using six different cut-offs. For each participant, the proportion of each node type was calculated by averaging across the six thresholds. For visualization purposes, individual node-type proportions were averaged across all the subjects to generate group averaged proportions of node types. A two-sample Mann-Whitney U test was used to compare these graph metrics between the ASD and TD groups, followed by FDR correction.

### 4.5) Node-role identification

To identify a specific global role of each brain region, the participation coefficient *(pc)* and within-module degree *(z)* of each node were categorized as a subtype of hub/non-hub for each subject, and then a node was regarded as a specific hub/non-non-hub based on its average frequency of occurrence in a group (i.e., the frequency of occurrence should be more than 50% of the subjects in the group). To avoid the arbitrariness of the network threshold, these measures were calculated for binarized and seven proportionally thresholded graphs.

To evaluate the reproducibility of functional cartography, we used the bootstrapping method. Within each group, these local measures were recalculated after bootstrapping, and then node roles were identified using resampled data. This procedure was repeated over 10,000 iterations followed by summing the occurrence frequency of a specific role of each node. We regard a node as a specific hub/non-hub based on a score that should be higher than the average frequency of occurrence.

## RESULTS

### 1. Global modular organization & ASD symptoms

We first examined three global modularity measures of brain networks using the Louvain algorithm (refer to Supplementary table 1 for Newman and Consensus agreement community detection results). The ASD group showed significantly reduced global modularity Q compared with the TD group (*p* = 0.001). We also observed a significant increase in the average number of modules (*p* = 0.02) and a significant decrease in the average modular size in the ASD group (*p* = 0.01) (**Table 2, Figure 1**). None of the whole-brain global measures of the modular organization were significantly associated with Autism Diagnostic Observation schedule (ADOS) behavioral scores (**Table 3**).

**Table 2.** Group differences in global modularity measures.

|  | ASD mean (± std) | TD mean(± std). | P<sub>value</sub> |
| --- | --- | --- | --- |
| <u>Whole-brain global modularity measures</u> |  |  |  |
| Global Modularity | 0.28 (±0.04) | 0.30(±0.03) | 0.001 |
| Average Number of modules | 2.99 (±0.52) | 2.81 (±0.39) | 0.02 |
| Average module size | 72.89 (±13.42) | 78.57 (±13.82) | 0.01 |
| <u>Whole-brain proportion of node types</u> |  |  |  |
| Connector hubs | 11.54(±2.58) | 10.77(±2.92) | 0.12 |
| Provincial hubs | 3.46(±2.45) | 4.43(±2.63) | 0.02 |
| Kinless hubs | 0.019(±0.05) | 0.003(±0.01) | 0.02 |
| Nonhub connector nodes | 19.09(±10.34) | 14.97(±8.04) | 0.008 |
| Peripheral nodes | 62.48(±10.14) | 66.59(±7.73) | 0.007 |
| Nonhub Kinless nodes | 0.002(±0.016) | 0.001(±0.007) | 0.65 |
| Ultra-peripheral nodes | 3.40(±2.62) | 3.24(±2.31) | 0.65 |

The group difference for global modularity measures was calculated using permutation test (10,000 iterations, $p < 0.05$ ) whereas whole-brain proportion of node types was calculated using Mann-Whitney U test.

**Table 3.** ASD Symptoms and global modularity measures.

|  | ADOS(Socio) | ADOS(RRB) | ADOS(total) | ADOS(severity) |
| --- | --- | --- | --- | --- |
| | $r_{\text{part.}}$ ($p_{\text{part.}}$) | $r_{\text{part.}}$ ($p_{\text{part.}}$) | $r_{\text{part.}}$ ($p_{\text{part.}}$) | $r_{\text{part.}}$ ($p_{\text{part.}}$) |
| <u>Whole-brain modularity measures</u> |  |  |  |  |
| Global modularity | -0.26 (0.05) | -0.08(0.55) | -0.25(0.06) | -0.18(0.17) |
| Average Number of modules | -0.09(0.49) | -0.006(0.96) | -0.08(0.55) | -0.08(0.52) |
| Average module size | 0.14(0.28) | -0.03(0.82) | 0.11(0.42) | 0.10(0.46) |
| <u>Whole-brain proportion of node types</u> |  |  |  |  |
| Connector hubs | 0.02 (0.83) | 0.09(0.51) | 0.06(0.65) | 0.06(0.65) |
| Provincial hubs | 0.01 (0.91) | -0.08(0.52) | -0.02(0.85) | 0.01(0.94) |
| Kinless hubs | -0.04 (0.73) | 0.18(0.18) | 0.03(0.77) | 0.0007 (0.99) |
| Non-hub connector nodes | -0.02 (0.83) | 0.13(0.32) | 0.03(0.80) | 0.03(0.77) |
| Peripheral nodes | 0.001 (0.99) | -0.12(0.38) | -0.05(0.71) | -0.03(0.79) |
| Nonhub Kinless nodes | 0.001 (0.98) | 0.09(0.47) | 0.04(0.75) | 0.05(0.68) |
| Ultra-peripheral nodes | 0.066 (0.63) | -0.08(0.55) | 0.02(0.88) | -0.08(0.54) |

ADOS, Autism Diagnostic Observation Schedule; RRB, Restricted and Repetitive Behaviors; Socio, Social-affective communication. The association between global modularity measures or whole-brain proportion of node types and ADOS behavioral scores were calculated using partial correlation while controlling for age, sex, site difference, and mean FD.

**Fig. 1:**
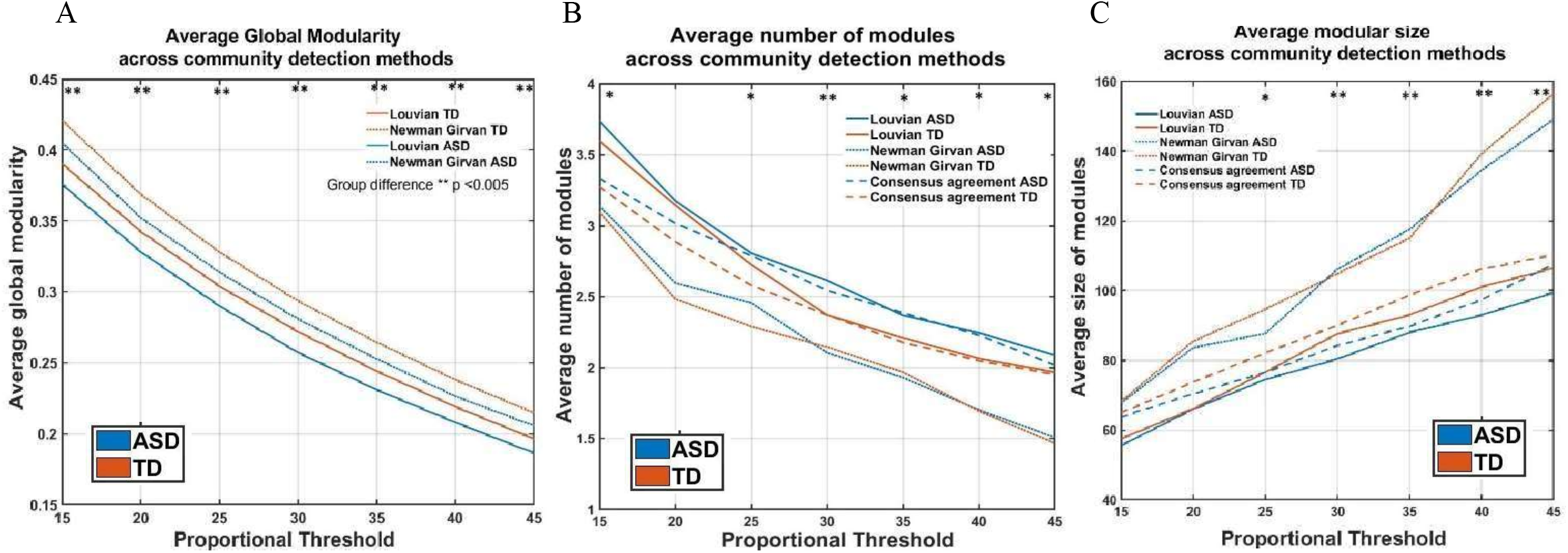
Group difference in global modularity measures across thresholds and community detection methods. There is significant reduction in global modularity (A), and the average modular size (C) in the ASD group as compared to TD population. There is significant increase in average number of communities (B) in the ASD participants. Graph metrics were calculated for proportionally threshold graph over a range of 15% to 45% with 5%increments. For each participant, these global measures were calculated by averaging across all the thresholds and for each group, these measures were averaged across all the subjects. The Louvain and Consensus agreement community-detection methods produced similar results as compared to Newman -Girvan methods (B & C).

### 2. Reduced similarity in modular composition in the ASD group

#### 2.1. Group-level analyses

A qualitative examination of group average modular structures (**Figure 2**) revealed that the overall network structure and functional connectivity were well preserved in both ASD and TD groups. The quantitative examination of the similarity between mean ASD and TD network structures (**Table 4)** using normalized mutual information (NMI) also showed a high level of similarity of network structures (averaged across proportional thresholds, mean NMI = 0.66, standard deviation = 0.13). (Refer to **Supplementary Table 2 & Figure 2** for group differences in modular composition across two atlas parcellations).

**Table 4.**
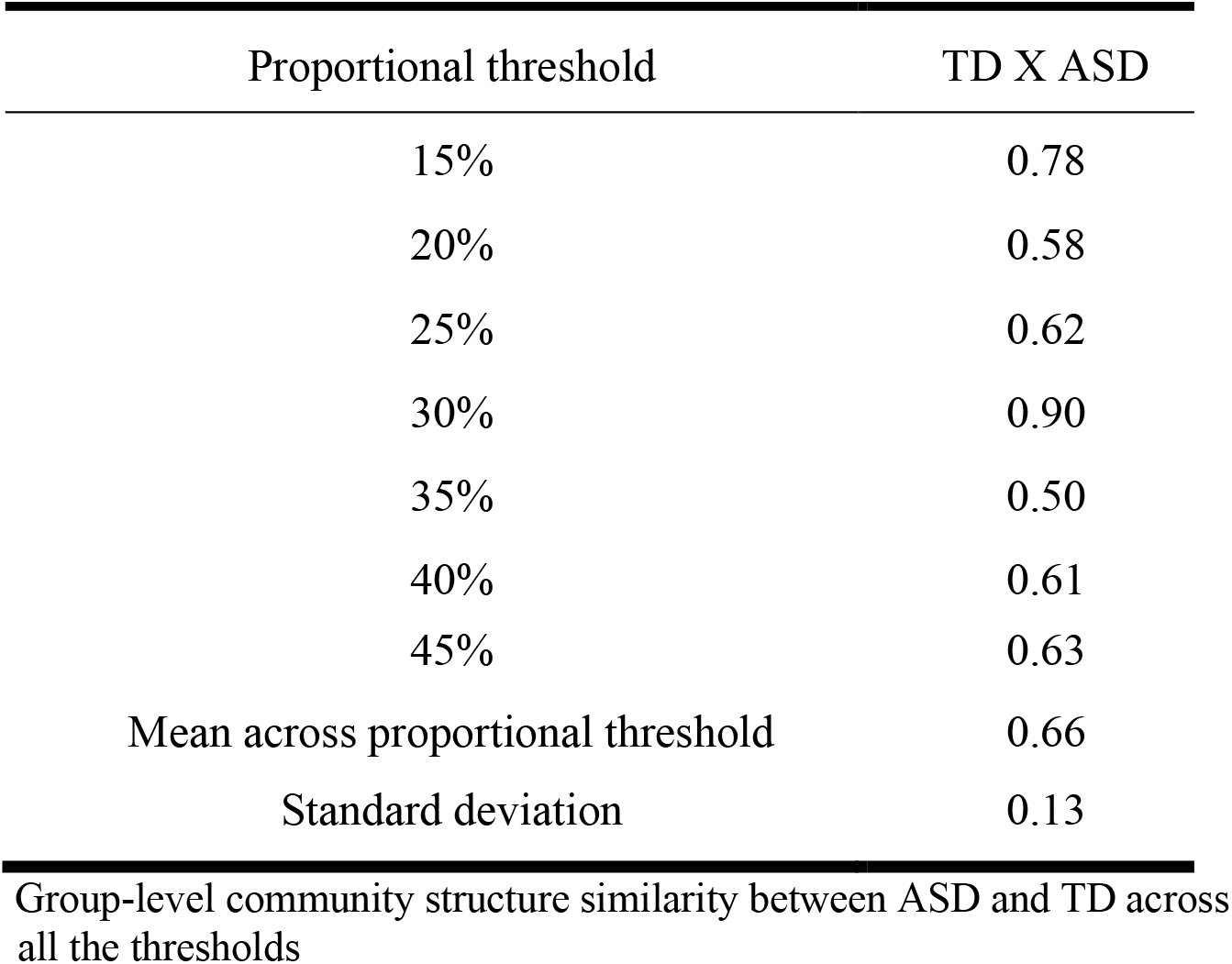
NMI values between ASD and TD (Supplementary)

**Fig. 2:**
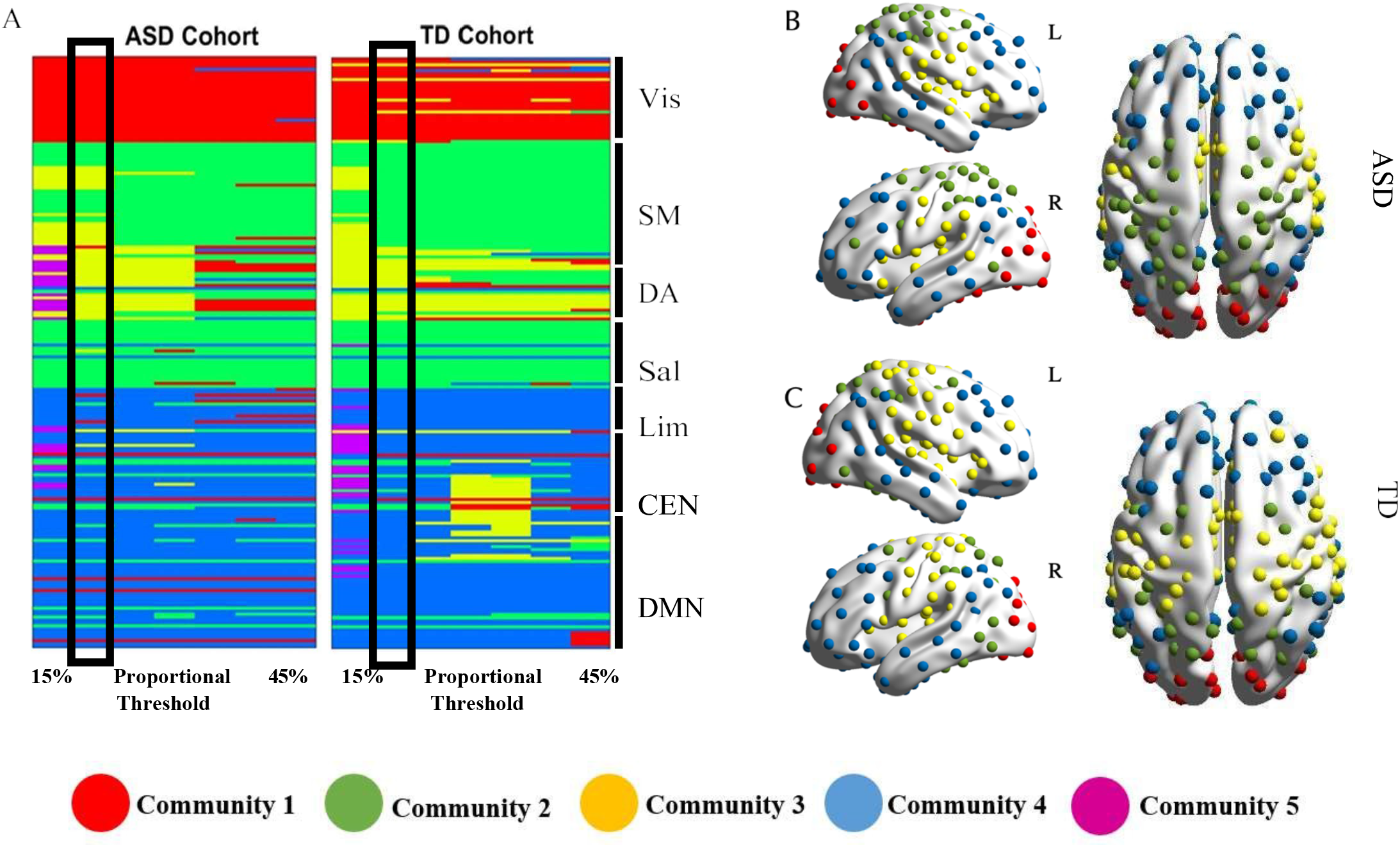
Group-level difference in the community structure revealed overall similarity between ASD and TD groups. A. The networks are on the vertical axis and graph densities are on the horizontal axis. Regions are colored by the community assignments of the ROIs. B. The community structure of ASD group at the 20% graph density. C. The community structure of TD group at the 20% graph density. The group-level community structures were detected using group-averaged functional connectivity matrices. The 20% graph density was used as representative community structure over a range of thresholds on the basis NMI.

#### 2.2. Subject-level analyses

Individual subject network partitions revealed that the mean NMI of all within-group subject pairs was significantly higher than the mean of all between-group subject pairs for all the thresholds except for one (**Table 5**). Additionally, across all the thresholds, the average within-group NMI for the ASD group was less than the average-within TD group NMI, reflecting the heterogeneity of network partitions in the ASD group. (Refer to **Supplementary Table 3, 4** for group differences in modular composition across two atlas parcellations).

**Table 5.** NMI permutation testing between ASD and TD group.

| Proportional Thresholds | P-value:<br>Real data><br>permuted data | Mean of all<br>within-group<br>NMI pairings | Mean of all<br>between-group<br>NMI pairings | Mean<br>within-TD NMI | Mean within-ASD NMI |
| --- | --- | --- | --- | --- | --- |
| 15% | 0.04 | 0.27 | 0.26 | 0.28 | 0.26 |
| 20% | 0.95 | 0.24 | 0.22 | 0.25 | 0.23 |
| 25% | 0.0002 | 0.21 | 0.20 | 0.22 | 0.21 |
| 30% | <0.0001 | 0.20 | 0.18 | 0.20 | 0.19 |
| 35% | <0.0001 | 0.18 | 0.16 | 0.19 | 0.17 |
| 40% | <0.0001 | 0.17 | 0.15 | 0.18 | 0.16 |
| 45% | <0.0001 | 0.16 | 0.14 | 0.17 | 0.15 |

Subject-level within group community structure similarity across all the proportional thresholds Permutation test with 10,000 iterations was performed with FDR correction.

#### 2.3. Node-level analyses

To further investigate the brain regions responsible for the significant group difference in the modular structure, we used follow-up permutation tests of community assignments (Alexander-Bloch et al., 2012). First, the significant nodes were selected using FDR corrected p<0.05 as a significance criterion, and the percent of significant nodes within each network was counted. **Figure 3A** shows that there was an inconsistency in community assignments in the ASD group with a high percentage of ROIs in the sensorimotor (SM) networks across all the threshold densities. Additionally, this community membership variability across groups was also contributed to by a smaller percentage of nodes in DMN, visual, salience, and executive control networks. To further illustrate the nodes contributing to the differences in the community structure and NMI between ASD and TD, the ROIs with significantly inconsistent/flexible community membership (FDR corrected, p<0.05) across all the threshold densities are represented on the brain (**Figure 3B**). The difference in the diagnostic network structure are driven by 38% of the nodes. Most notable community membership discrepancies were contributed by lateral occipital cortex (LoCc), precentral gyrus (PrG), postcentral gyrus (PoG), superior temporal gyrus (STG), dorso-medial prefrontal cortex (dmPFC), precuneus (Pcun), inferior parietal lobule (IPL), and lateral prefrontal cortex (lPFC).

**Fig. 3:**
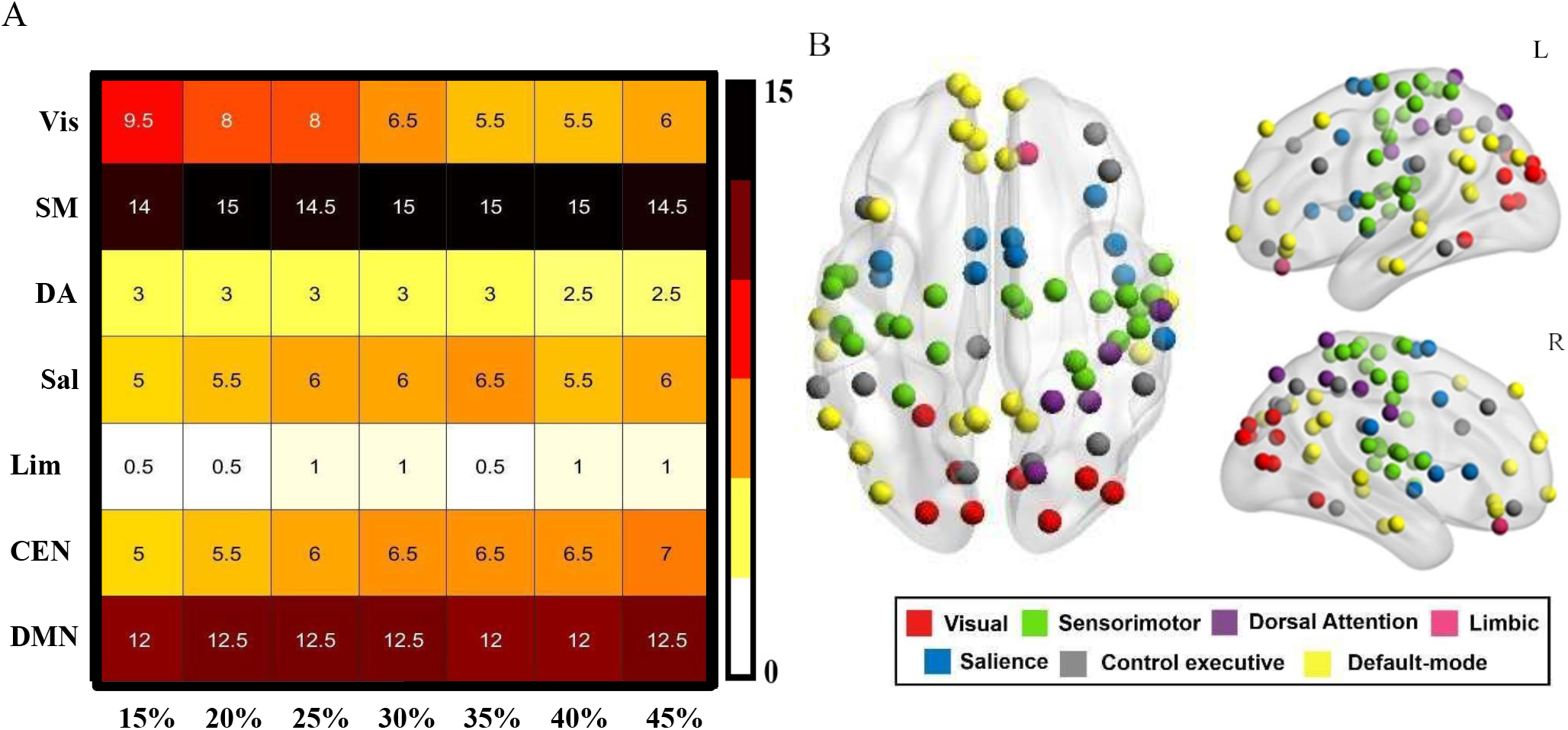
Subject and Node-level community structure differences between ASD and TD. A. Subject-level Phi test results revealed that nodes of sensorimotor (SM), Default-mode (DMN) and Visual (Vis) networks, significantly altered their community assignments. Dark colors in the map represents higher fraction of nodes with significant (FDR corrected, p< 0.05) alterations in community assignments at that graph density. B. Node-level Phi test results exhibited 38% of nodes with significant (FDR corrected, p< 0.05) alteration in community assignments. ROIs from sensorimotor, default-mode, salience and visual networks revealed significant inconsistency in modular membership across all the graph densities. ROI color represent their respective brain functionalnetwork.

#### 2.4. Association with ASD symptoms

The community assignment consistency values (Pearson’s phi coefficient) of the majority of inconsistent nodes (dmPFC, PCun, LocC, medial ventral occipital cortex (MvocC), lPFC, cingulate gyrus (CG)) were negatively associated with ADOS-social, total and severity scores whereas these community assignment consistency values of nodes such as (dmPFC, PoG, lPFC, LocC, MvocC) were also negatively associated with restricted and repetitive behavior (RRB) scores. These results suggest that with increasing symptom severity the tendency of nodes to switch modules within the ASD group also increases (i.e., reduced stability of nodes within their respective modules).

### 3. Altered modular connectivity

To investigate group differences in modular connectivity, we used two measures: participation coefficient PC (inter-modular connectivity) and within-module degree Z (intramodular connectivity). In ASD, a significantly (FDR corrected, p<0.01) increased participation coefficient was observed for nodes in bilateral PCun, bilateral PrG, and PoG, rIPL, and rPCun (**Table 6, Figure 4A, B**). The nodes with significant differences in within-module degree did not survive FDR correction, however, in ASD significant (all *p*<0.05) increase in intramodular connectivity was contributed to by nodes of bilateral LoCc, bilateral temporal lobe, and left (medial orbito-frontal gyrus) mOFG. A significant decrease in within-module connectivity was observed in nodes of right IFG and SFG, bilateral paracentral lobules (PCL), bilateral PoG, rIPL and frontal eye field (FEF), and left lingual cortex and right cuneus cortex consisting of the MvocC node (**Table 6, Figure 4C, D**). The within-module connectivity values of bilateral nodes of dmPFC were negatively (*r* = -0.3, *p* = 0.01) associated with the ADOS RRB score whereas the between-module connectivity of the same nodes was positively correlated (*r* = 0.3, *p* = 0.02) with ADOS severity scores. The average participation coefficient scores of DMN, salience, and sensorimotor networks were also negatively correlated with global modularity (*r* = -0.58, *p*<0.0001) thus further suggesting that the increased between-module connectivity in ASD results in less robust modular organization contributed by nodes of these networks. Furthermore, in the ASD group, there was a strong negative correlation between the participation coefficient with community assignment consistency scores of the nodes with increased flexibility (which switched their modules across different subjects) (**Figure 4E, F**). (Refer to **Supplementary Figure 1** for group-differences in modular connectivity across two atlas parcellations).

**Table 6.** Modular connectivity.

| Brain regions | Network | Hem | x | y | z |
| --- | --- | --- | --- | --- | --- |
| Between-Module connectivity |  |  |  |  |  |
| ASD>TD |  |  |  |  |  |
| PrG<sup>a</sup> | SM | L | -47 | -9 | 46 |
| PoG<sup>a</sup> | DA | L | -41 | -35 | 47 |
| lITG<sup>a</sup> | DMN | L | -60 | -19 | -22 |
| vPCun<sup>a</sup> | DMN | L | -5 | -55 | 27 |
| lingual MVocC<sup>a</sup> | Vis | R | 16 | -46 | -1 |
| PrG<sup>a</sup> | SM | R | 59 | 0 | 10 |
| PoG<sup>a</sup> | SM | R | 56 | -11 | 14 |
| vIPL<sup>a</sup> | DA | R | 59 | -16 | 34 |
| PrCv | Sal | R | 51 | 4 | 40 |
| MTG<sup>a</sup> | DMN | R | 61 | -13 | -21 |
| PCun<sup>a</sup> | DMN | R | 7 | -49 | 31 |
| TD>ASD |  |  |  |  |  |
| mSTG | Lim | R | 47 | -12 | -35 |
| Within-Module connectivity |  |  |  |  |  |
| ASD>TD |  |  |  |  |  |
| PCL | Sal | L | -11 | -35 | 46 |
| TD>ASD |  |  |  |  |  |
| infLOcC | Vis | L | -27 | -95 | -12 |
| PhG | Lim | L | -29 | -6 | -39 |
| mSupLOcC | Vis | R | 16 | -85 | 39 |

Mann-Whitney U test over 10,000 iterations; $p < 0.01$ ; \*FDR correction survived
Hem, Hemisphere; L, left; R, right; xyz, Montreal Neurological Institute template brain (MNI) coordinates Vis: Visual, SM: Sensorimotor, DA: Dorsal-attention, Sal: Salience, Lim: Limbic, DMN: Default-mode.

**Fig. 4:**
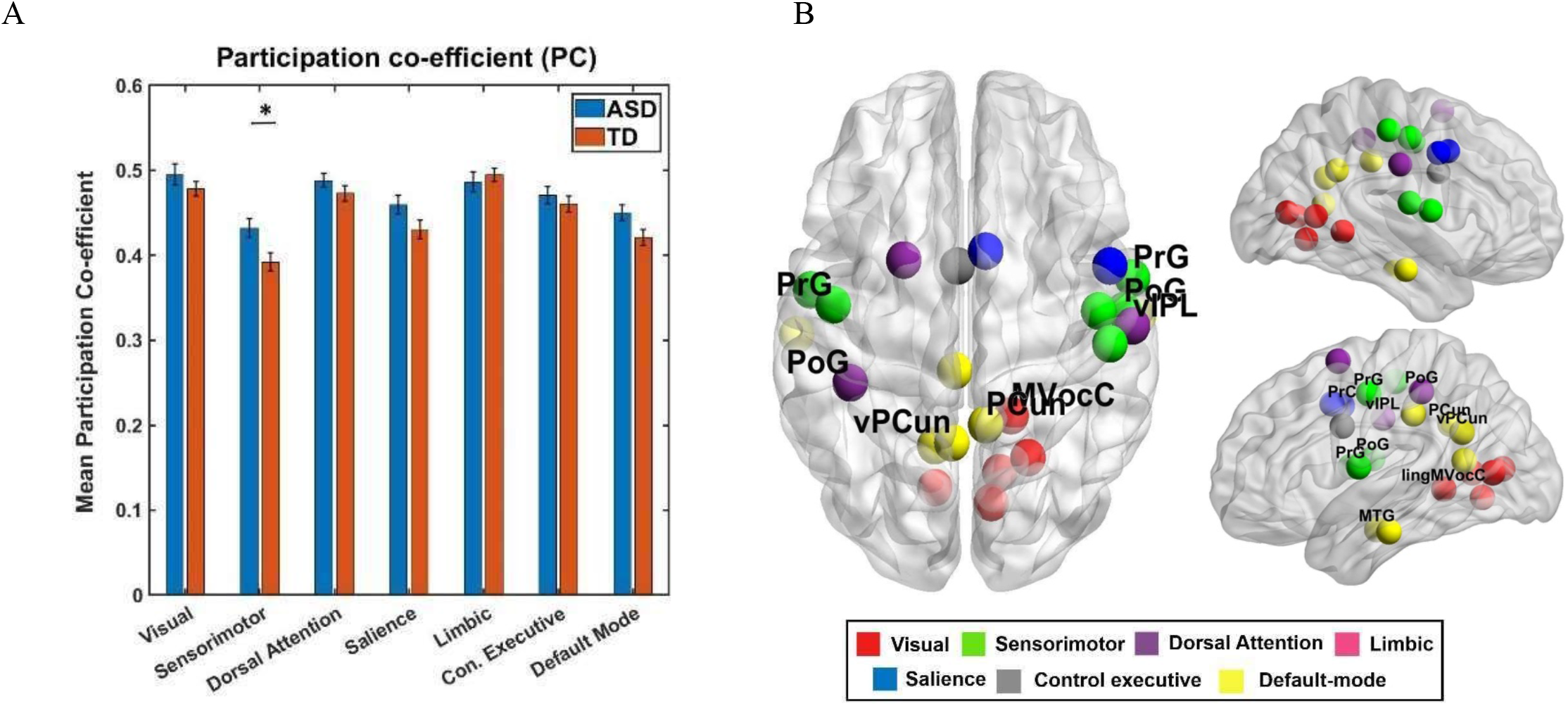
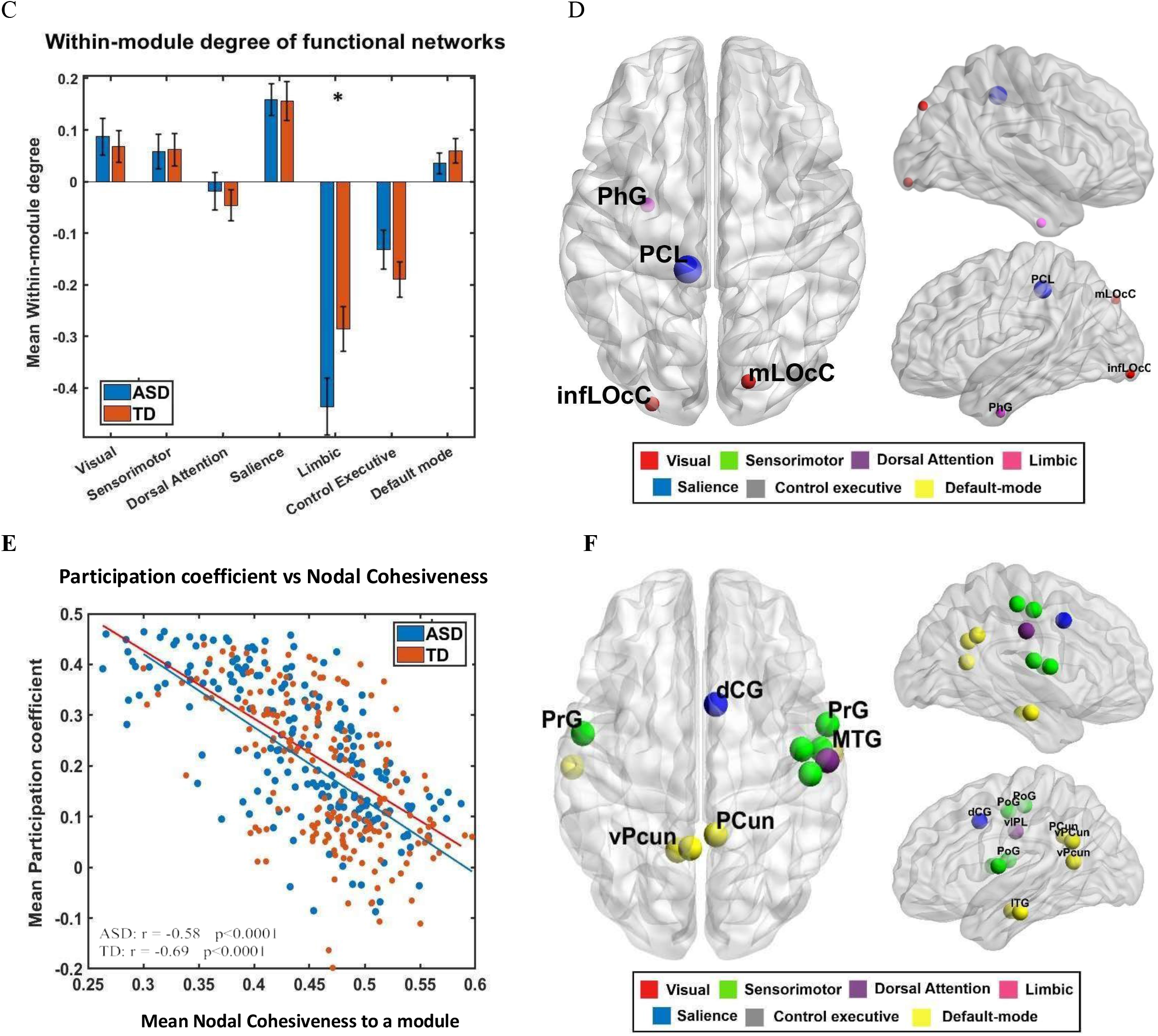
Group difference in modular connectivity and association between inter-modular connectivity & modular assignment inconsistency. A. The network- level between–module connectivity was calculated by averaging the PC scores of all the nodes within each network. Sensorimotor network has significantly (FDR corrected, p<0.05) higher between-module connectivity in ASD group as compared to TD group. B. The node-level results exhibited that the ROIs of Default-mode, sensorimotor, salience and visual network have significantly (FDR corrected, p<0.05) increased inter-modular connectivity in ASD group. C. The network-level within-module connectivity revealed that limbic network has significantly (FDR uncorrected, p<0.05) reduced intra-modular connectivity in ASD group. D. The node-level intra-modular connectivity analysis showed mixed results. Graph metrics were calculated for binarized and proportionally thresholded graphs (15% to 5% with 5% increments). For each participant, these graph metrics were calculated by averaging across all the thresholds and for each group, these measures were averaged across all the subjects. E. There is a significant negative correlation between average participation coefficient and average nodal cohesiveness to a module (consistency of modular membership) for both the groups – ASD and TD. F. The nodes reflecting significant negative association between average PC scores and nodal cohesiveness are from Default-mode, sensorimotor, salience and Dorsal attention network.

### 4. Functional cartography reveals significant differences in node types in the ASD group

The topological roles of ROIs in facilitating within and between module communication were characterized using participation coefficient (PC) and within-module degree Z. **Figure 5B** shows that only 15.18% of nodes were characterized as hubs (i.e., kinless, connector, or provincial hubs) and the majority of the node types were peripheral and non-hub connectors. In ASD, proportions of peripheral and provincial nodes were significantly reduced, whereas proportions of non-hub connector nodes and connector hubs were significantly increased (**Table 2**). However, the proportion of node types showed no significant association with ADOS scores (**Table 3**) (Refer to **Supplementary Figure 3** for group-differences in modular composition across two atlas parcellations).

**Fig. 5:**
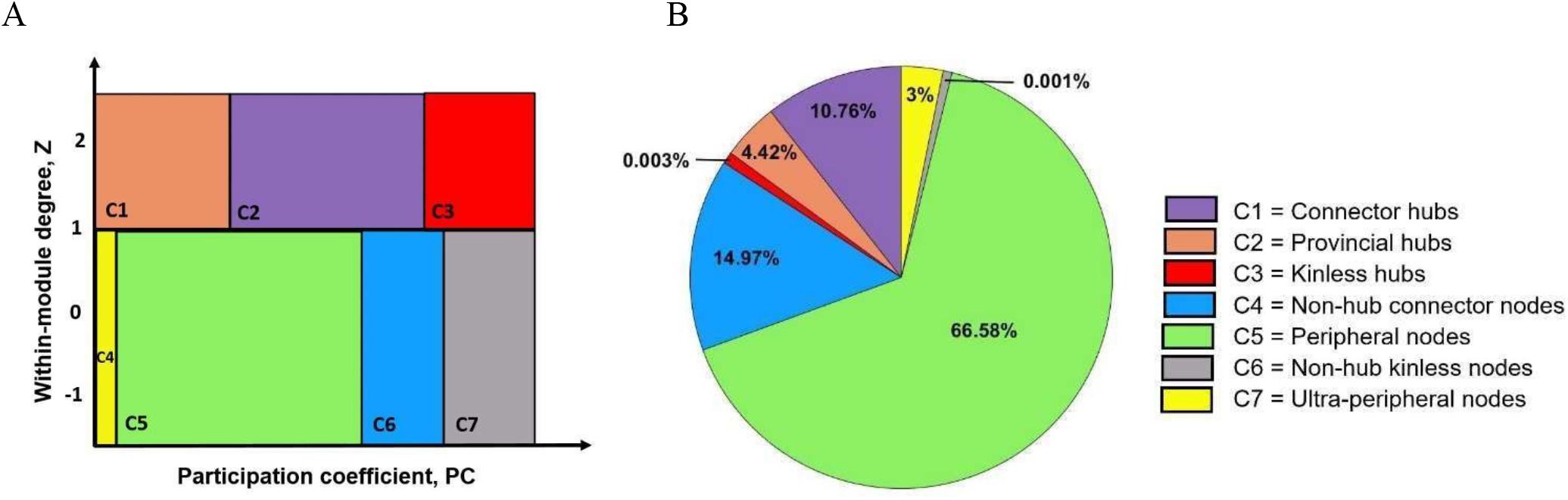
Functional cartography exhibited altered proportion of different node types in ASD group. A. Nodes classified into 7 types based on participation coefficient, PC and within-module degree, Z (Guimerà and Amaral 2005). B. Whole-brain group-averaged proportions of node types. This was calculated separately on binarized and proportionally thresholded graphs (15%, 20%, 25%, 30%, 35%, 40%, and 45%). For each subject, individual node-type proportion was calculated by averaging across the six thresholds

### 5. Nodal-role identification

We further examined brain regions which contributed to the altered proportions of the node types. Qualitative examination, analyzed using the group average matrix, shows that the majority of peripheral nodes in TD converted to non-hub connectors (PrG, PoG, lPFC, PCun, ventral PFC, insula) and few peripheral nodes to connector hubs in ASD (bilateral dmPFC, lPFC, right PoG). Whereas the provincial nodes in TD switched their roles to connector nodes in ASD (like the left IPL from DMN (**Figure 6A, B**)). Subject-level analyses revealed that the ASD group had 19 non-hub connector nodes and 5 connector hubs as compared with the TD group with 4 non-hub connector nodes and 2 connector nodes (**Figure 6E, F, G, Table 7, 8**). Common non-hub connectors (8 nodes) were from the left LoCc, left FuG, bilateral PCL, and rOFC. Additionally, the ASD group had 13 non-hub connectors from the bilateral temporal pole and FEF, left OFC and right medial temporal lobe, and bilateral vPFC. Common connector hubs were from bilateral STG, bilateral Insula cortex, and right dmPFC. Additionally, the ASD group had 5 connector hubs from bilateral STG, bilateral posterior cingulate cortex (PCC), and left FuG, whereas the TD group had 2 extra connectors from the dmPFC.

**Table 7.** A list of connector hubs.

| Brain regions | Network | Hem | x | y | z |
| --- | --- | --- | --- | --- | --- |
| **ASD-specific hubs** |  |  |  |  |  |
| Lingual MVocC | Vis | L | -10 | -67 | -4 |
| STS | SM | L | -53 | -24 | 9 |
| STS | SM | R | 51 | -15 | 5 |
| dCG | Sal | R | 7 | 9 | 41 |
| **TD-specific hubs** |  |  |  |  |  |
| mOFG (dmPFC) | DMN | L | -12 | 63 | -6 |
| mSFG (dmPFC) | DMN | L | -8 | 59 | 21 |
| **Common hubs** |  |  |  |  |  |
| dINS | Sal | R | 46 | -4 | -4 |
| dmPFC | DMN | R | 8 | 58 | 18 |
| STG | SM | L | -51 | -4 | -2 |

Bootstrap method with FDR correction ( $p < 0.001$ )
Hem, Hemisphere; L, left; R, right; xyz, Montreal Neurological Institute template brain (MNI) coordinates Vis: Visual, SM: Sensorimotor, DA: Dorsal-attention, Sal: Salience, Lim: Limbic, DMN: Default-mode.

**Table 8.**
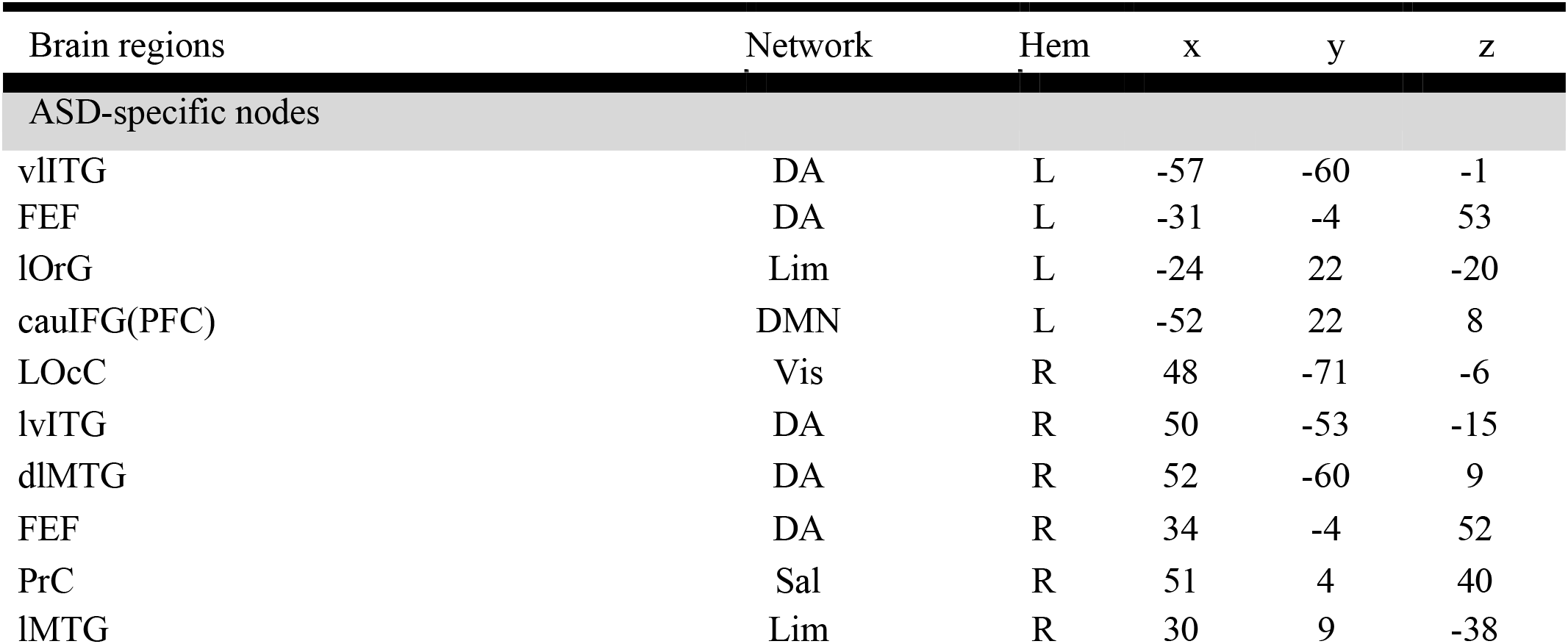
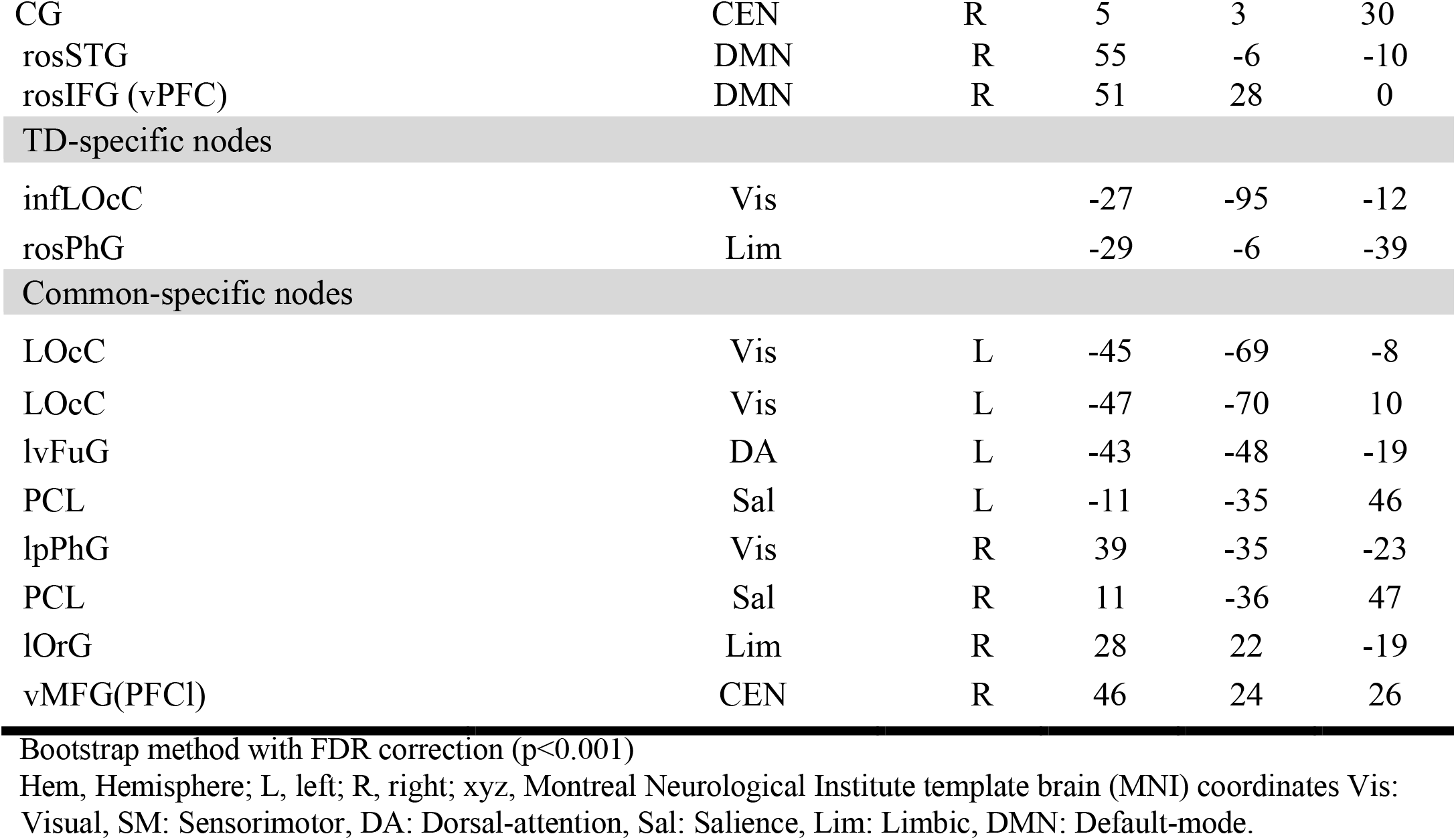
A list of non-hub connector nodes.

**Fig. 6:**
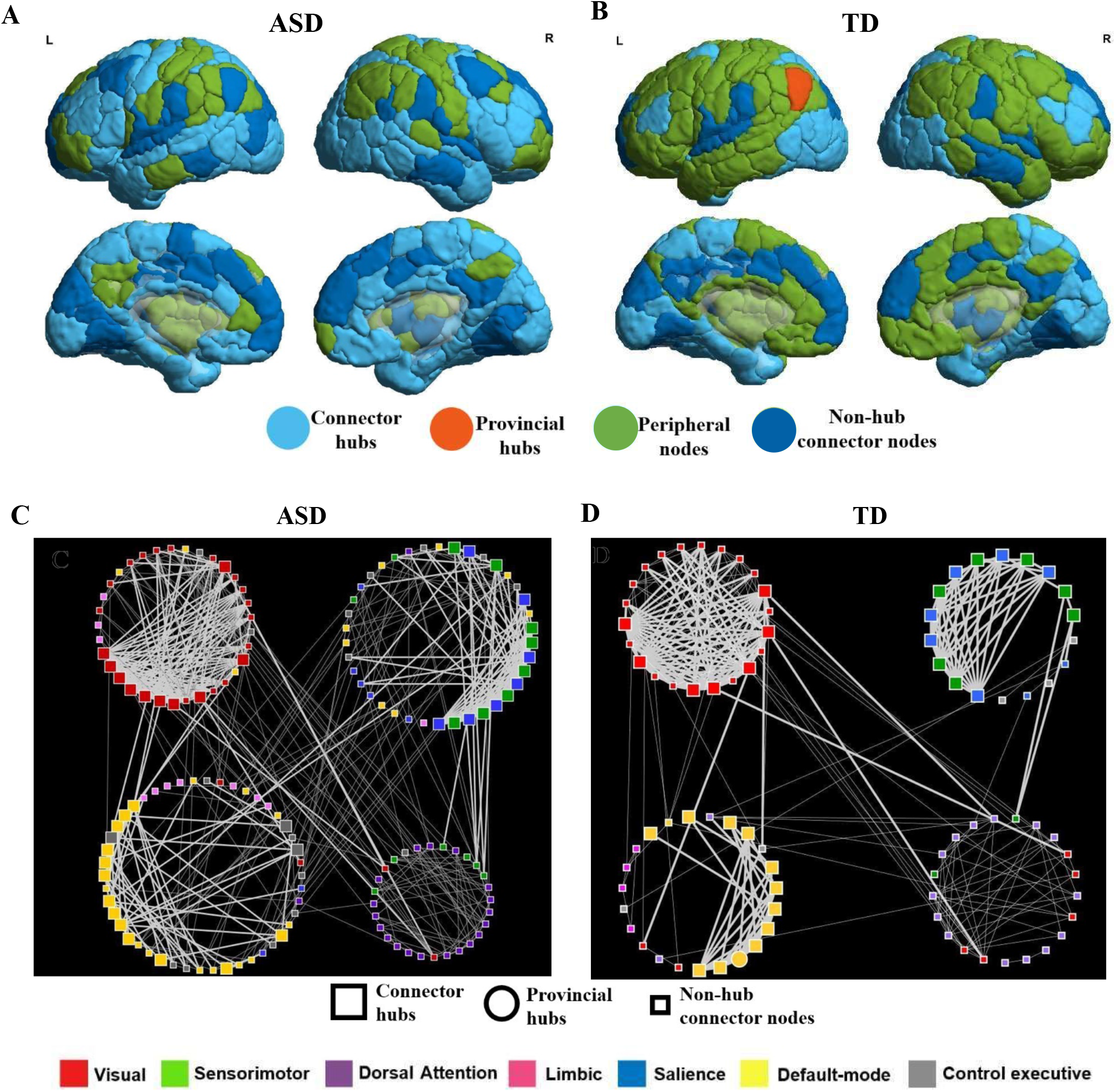
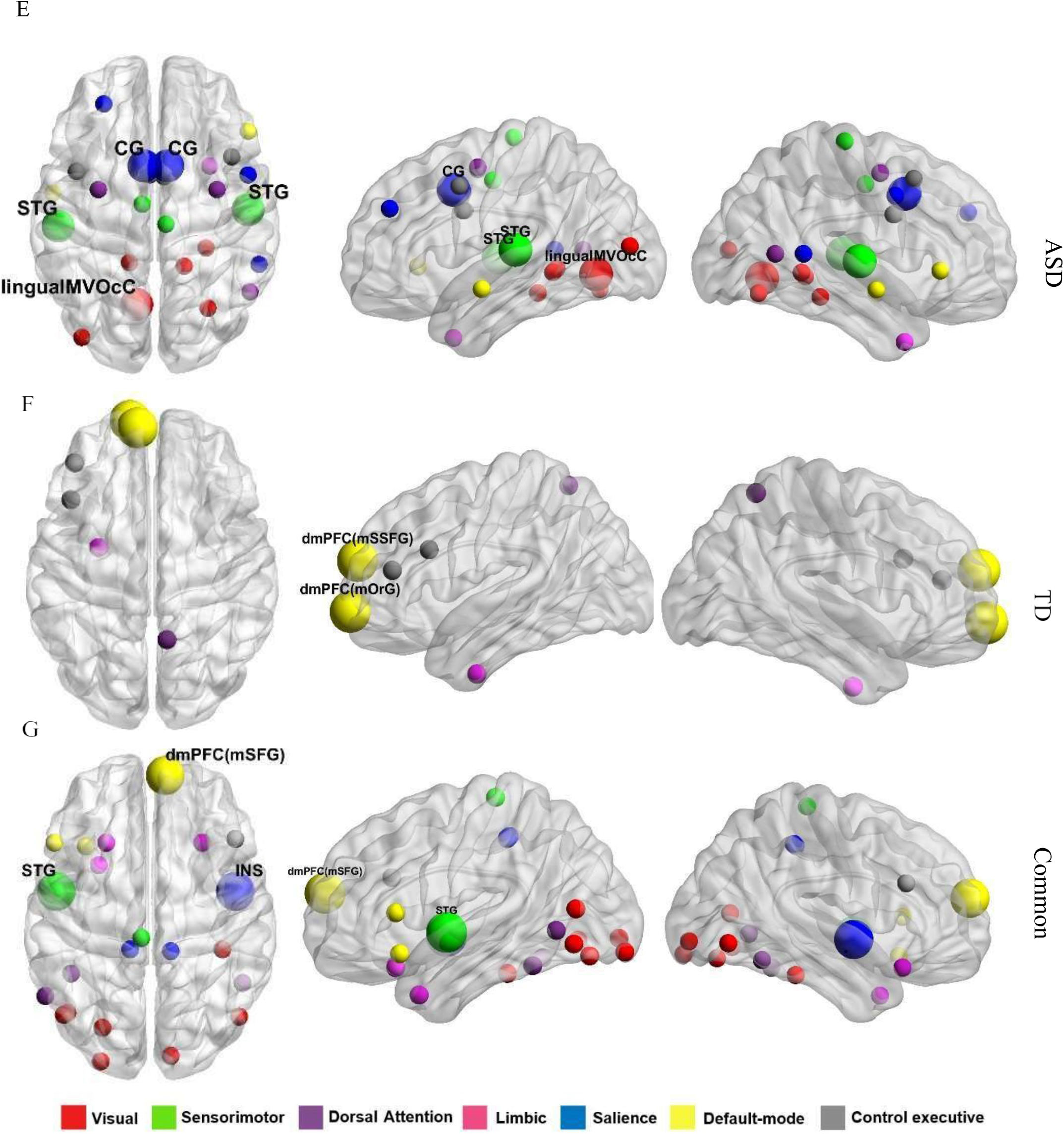
Group-differences in topological roles of nodes. A. The group-level node-roles identification analysis revealed that overall there are more connector hubs and non-hub connector nodes in ASD group in temporal, and frontal areas B. At group-level, there are more peripheral nodes and provincial hubs in parietal, frontal and temporal areas whereas the ratio of connector hubs and non-hub connectors is limited to occipital cortex, medial temporal and parietal areas. Figure C & D represent the four communities with the distribution of different-node types across seven functional networks. The bigger squares are connector hubs and smaller square are non-hub connector nodes. Figure E, and F represent significant connector hubs (bigger sphere) and non-hub connector nodes (smaller sphere) in ASD and TD group respectively. G. There are few significant connector hubs and non-hub connector nodes which are common across both thegroups.

The majority of non-hub connector nodes in the ASD group had increased between-module connectivity (FDR uncorrected, all *p*<0.05), thus reflecting the influence of these nodes as non-hub connectors on increased inter-module connectivity in ASD populations (**Figure 6C**). Similarly, connector hubs (left FuG and rPCC) in ASD had increased inter-modular connectivity (FDR uncorrected, all *p*<0.05).

## DISCUSSION

Autism spectrum disorder is a ‘critical period’ neurodevelopmental disorder associated with atypical development of brain networks. Globally atypical functional brain networks may arise from the increased ratio of excitation/inhibition balance resulting in hyperexcitability and thus cortical instability in ASD (Rubenstein and Merzenich 2003). Furthermore, these functional brain network atypicality in ASD may also be associated with synaptic cell-adhesion molecules and altered expressions of genes encoding these molecules, which play important roles in synaptic formation and axonal guidance (Arons et al., 2012; Földy et al., 2013; Jamain et al., 2003; Redies et al., 2012). Most of the neuroimaging studies examining brain network organization in ASD have focused on older children, adolescents, and adults, with less work on younger children (5-10 years). In our study, we systematically investigated global and regional module properties related to atypical cognitive skills and social communication deficits in younger children with ASD.

We used graph-theoretical analysis applied to resting-state fMRI data from ABIDE. We report evidence for alteration of local node properties on the atypical modular organization of functional brain networks in children with ASD. Children with ASD show reduced global modularity as well as within-group modular composition consistency compared with age-matched TD children. These results were also significantly associated with alterations in modular connectivity in ASD. Further investigations of the underlying nodal cause of the global modular organization revealed that the alterations in roles of nodes from sensorimotor, salience and, default mode networks resulted in alteration of modular connectivity, thereby affecting the modular composition and global modularity in ASD.

### Alteration in global modularity in ASD

As reported in the previous literature, we observed a reduction in global modularity that might reflect hyper-synchronized brain networks (i.e., fewer connections within modules and more connections between modules) in ASD (Harlalka et al. 2018; Henry, Dichter, and Gates 2018; Keown et al. 2017; Rudie et al. 2013). Additionally, we also observed an increase in the number of modules, along with a significant decrease in the modular size in the ASD group, which is indicative of the partitioning of brain networks into modules of smaller sizes in ASD participants compared with TD. These alterations in global modular properties might be reflective of the apparent randomness of functional brain networks in ASD individuals (Henry, Dichter, and Gates 2018; Keown et al. 2017; Rudie et al. 2013) and might result from alterations in graph-theoretical properties such as reduction in clustering coefficient and characteristic path length as reported in several previous studies of adults with ASD (Itahashi et al. 2014).

### Reduced similarity in modular composition in ASD

Reconfiguration of brain networks might play an integral role in executing cognitive functions. Previous studies report that alterations in modular reconfiguration might be associated with reduced cognitive functioning with age (Meunier et al. 2009) and clinical disorders such as schizophrenia (Heuvel et al. 2010). Group-level modular structure composition analysis reflects that both groups shared nearly equal amounts of information, suggesting that at the gross level, functional brain organization is relatively well-preserved in ASD compared with TD.

However, subject-level analyses revealed that significant group differences in community composition reflected an increased heterogeneity (i.e., reduced similarity) in modular composition within ASD compared with the TD group. Keown 2017 also showed this reduction in community structure similarity in individuals with ASD. Our study also revealed that global levels of modular reconfiguration and heterogeneity in modular structure in ASD might result from reduced cohesiveness of nodes to their modules across subjects, especially nodes of networks such as visual, sensorimotor, executive control, and DMN.

### Modular cohesiveness of nodes directed by their modular connectivity

Alteration in modular connectivity is essential for the integration and segregation of networks, which permits a reconfiguration of the global modular organizations to perform different types of cognitive tasks such as working memory or tapping fingers (Stanley et al. 2014; H. E. Stevens et al. 2017; Cohen and D’Esposito 2016). Modular connectivity alters the flexibility of the nodes to switch modules to mediate the exchange of information across modules (Harlalka et al. 2019). Our findings show that the increase in between-modular connectivity of nodes belonging to three functional brain networks, namely DMN, SMN, and salience, was negatively associated with their modular cohesiveness. This provides an explanation that these nodes with increased between-module connectivity (BMC) interact more with other modules and thus possibly switch their modules more frequently in the ASD group compared with the TD group. These findings validate that the altered inter-modular connectivity of DMN, SM, and salience networks not only impacts the overall modular composition but also influences global modularity. Furthermore, this overall increase in BMC along with a reduction in global modularity in the ASD group reflects that the system exhibits less robust modular organization and explains the possible atypical synchronicity of the functional brain networks reported in ASD individuals.

### Nodal roles altered in ASD

The brain regions/nodes with different functional roles influence the flow of information within and between modules (Luo and Constable 2022) and thus the proportion of different types of nodes will further influence the global flow of information, efficient integration /segregation of functional brain networks, and thus also affect cognitive performance (M. A. Bertolero et al., 2017; Maxwell A. Bertolero et al., 2018). Thus, the proportions of nodes with different topological roles are also considered global properties of modular network organization (Van den Heuvel & Sporns, 2013). To facilitate information flow across and within the modules, different types of nodes have differing abilities to switch modules, thus different node types also have different modular membership consistency (e.g., connector hubs have more flexibility to switch modules and play a crucial role in exchanging information within and across modules).

Our results demonstrate that children with ASD had relatively higher proportions of non-hub connectors (NHC) (responsible for between-module connectivity) and reduced proportions of peripheral non-hubs (NHP) and provincial hubs (HP) (maintain more within-module connectivity) compared with TD children. The follow-up analysis revealed that peripheral nodes in TD converted to non-hub connectors in ASD, whereas provincial hubs converted to connector hubs in ASD. These findings explain the observed increase in between-module connectivity in ASD due to increased proportion of NHC nodes and decreased within-module connectivity due to decreased proportions of HP and NHP.

As reported in previous literature, major common connector hubs were from sensorimotor, salience, and default mode networks, whereas children with ASD had more connectors from sensorimotor and salience networks (Siddharth Ray et.al.2014, Keown 2017). These nodes being connectors also had higher flexibility to switch modules and result in increased between-modular connectivity. Furthermore, another node type exhibiting increased modular connectivity in ASD was non-connector nodes, from default mode network and dorsal attention and limbic networks. These results reflect that major large-scale network-level alterations in children with ASD involve brain regions of the DMN, SMN, and salience networks (Lynch et al., 2013; Marshall et al., 2020; Uddin et al., 2013, 2015; Uddin & Menon, 2009).

## LIMITATIONS

The study was based on multi-site data from ABIDE I and II, thus, to eliminate multi-site covariations we used statistical tests. However, co-variations caused by differing scanning protocols and scanners were not considered. Furthermore, since ASD is more prevalent in males and to avoid co-variations resulting from effects of gender, we focused on male individuals with ASD. Further investigation will be required to study gender effects on modular properties of functional brain networks in ASD. This study had a small sample size thus, the current findings await replication with a larger sample size using data from sites other than ABIDE I and II.

## ACKNOWLEDGMENTS AND DISCLOSURES

This study was supported by NBRC Core funds and Computing Facility, Ramalinga swami Fellowships (Department of Biotechnology, Government of India) to DR (BT/RLF/Re-entry/07/2014). DR was also supported by SR/CSRI/21/2016 extramural grant from the Department of Science and Technology (DST) Ministry of Science and Technology, Government of India. DR and AB acknowledge the generous support of the NBRC Flagship program BT/ MEDIII/ NBRC/ Flagship/ Program/ 2019: Comparative mapping of common mental disorders (CMD) over the lifespan.

## Funding Resources

NBRC core, Department of Biotechnology, Govt. of India (Grant Nos: BT/RLF/Re-entry/07/2014) Department of Science and Technology, Ministry of Science and Technology (Grant No. SR/CSRI/21/2016), NBRC Flagship program BT/ MEDIII/ NBRC/ Flagship/ Program/ 2019.

## FINANCIAL DISCLOSURES

The authors state that there are no biomedical financial interests or potential conflicts of interest, regarding the work presented here, by any of the authors.

## Glossary

BOLD: Blood oxygen level dependent
FDR: False-positive discovery rate
FD: Frame wise displacement (head motion)
FIQ: Full Intelligence quotient (verbal + non-verbal)
ADOS: Autism Diagnostic Observation schedule
RRB: Repetitive and restriction behavior
R-fMRI: Resting state functional magnetic resonance imaging
SFC: Static Functional connectivity
ASD: Autism spectrum disorder
TD: Typically developing
ADHD: Attention deficit and Hyperactivity disorder
ABIDE: Autism Brain imaging data exchange
DPARSF: Data processing assistant for resting-state fMRI.
MPRAGE: Magnetization Prepared Rapid Acquisition Gradient Echo
DARTEL: A Fast Diffeomorphic Registration Algorithm
FWHM: Full width at half maximum
ROI: Region of Interest
NMI: Normalized Mutual Information

## Supplementary material

### METHODS

#### 1. Threshold selection

Several studies suggest that graph properties like modularity are strongly influenced by the different graph densities (Ginestet, Nichols, Bullmore, & Simmons, 2011). Thresholding a graph based on connection strength can yield differences in graph density, thus influencing graph properties and creating a biased comparison of graph metrics between the groups (Bullmore & Bassett, 2011a; Ginestet et al., 2011; Schwarz & McGonigle, 2011; van Wijk, Stam, & Daffertshofer, 2010). Thus, to avoid biases due to inter-subject differences in graph density, we equalized network density between subjects by thresholding graphs to retain an equal proportion of the strongest positive connections (negative connections were excluded), followed by binarizing (recommended to retain individual differences in network topology graphs (Bullmore & Bassett, 2011b; van Wijk et al., 2010). We thresholded our correlation matrix over a range of thresholds, retaining the strongest 15, 20, 25, 30, 35, 40, and 45% edges. This resulted in seven thresholded graphs for each subject. The upper thresholded 45% was chosen because at the 45% threshold, and the weakest edge corresponds to a statistically significant (p<0.05) minimum correlation coefficient (0.2) across 170 functional images. At the lowest threshold, networks become fragmented, thus breaking down network properties. Graph density with 15% of strongest edges was chosen as the lower threshold (significant weakest connections r =0.15, p-value < 0.05) at which the graph was fully connected for each subject. We tested the group differences in modularity and community structure at all connection densities

#### 2. Modular composition analyses

##### 2.1) Group-level modularity composition analyses

We constructed a group representative modularity partition by applying the Louvain method to an unthresholded group averaged functional connectivity matrix after excluding all the negative weights.

However, given the controversy regarding comparing modularity partitions of group averaged matrices with each individual’s modular partition (Simpson, Moussa, & Laurienti, 2012), using the NMI approach, we compared the similarity of our representative modularity partition with each individual’s modularity partition (Meilă, 2007). Furthermore, to assess whether the similarity between the modular partition of representative group averaged matrices and individual matrices were because of chance or group membership, we checked the similarity of our representative modularity partition with randomized individual matrices. Random networks were created using the MATLAB-based Brain Connectivity toolbox function: randmio_dir_connected.m, which preserves the degree of connectedness of each node in the true network. However, since this measure is graph density sensitive, we repeated this computation with all seven proportionally thresholded graphs (15% to 45% with an increment of 5%) to obtain an NMI value of similarity between community assignments for each density graph (Lerman-Sinkoff & Barch, 2016).

##### 2.2) Subject-level modularity composition analyses

Using the NMI approach, we can study the difference in community structure within and between groups at the individual subject level and explore within-group heterogeneity of network structure. The community structure variance can be more reliably explained by group membership than by chance only if the average NMI between all the pairs of subjects within a group is higher than the average NMI between pairs of subjects selected at random. The group label permutation method developed by (Alexander-Bloch et al., 2012) was used to determine the statistical similarity of subjects within a group and across the two groups. For all six proportionally thresholded graphs, the true within-group NMI average and true between-group NMI average were calculated by taking the mean of NMI values computed for each pair of participants. Over 10,000 iterations, group labels were randomly permuted, and mean within-group NMI was calculated for permuted data at each density threshold. Further, at each density thresholded, the p-value was calculated as the ratio of the total number of instances when the actual mean within-group NMI is less than the permuted mean within-group NMI, relative to the number of permutations (Lerman-Sinkoff & Barch, 2016).

The above permutation test does not reveal how the community affiliation/assignment of a particular ROI may differ across the ASD and TD groups. Using the second permutation test developed by (Alexander-Bloch et al., 2012), we assessed node-specific statistical differences in community structure between groups. Pearson’s phi coefficient (-1<ϕ<1) (Pearson,1990) approach was used to quantify node similarity between two subjects in terms of a node’s functional community. If there is a difference in a node’s community between two groups, then the averaged between-group phi coefficient value should be lower than the averaged within-group phi coefficient value. Permutation procedures can be used to generate a p-value comparing the average of within-group Phi values in real data and randomized/permuted data. Thus, for each density threshold, a set of p-values was generated indicating whether a given ROI’s community was more similar for subjects within the same group than in randomized groups. This test was performed at each density threshold, for each node over 10,000 iterations, followed by FDR correction.

**Supplementary Table 1:**
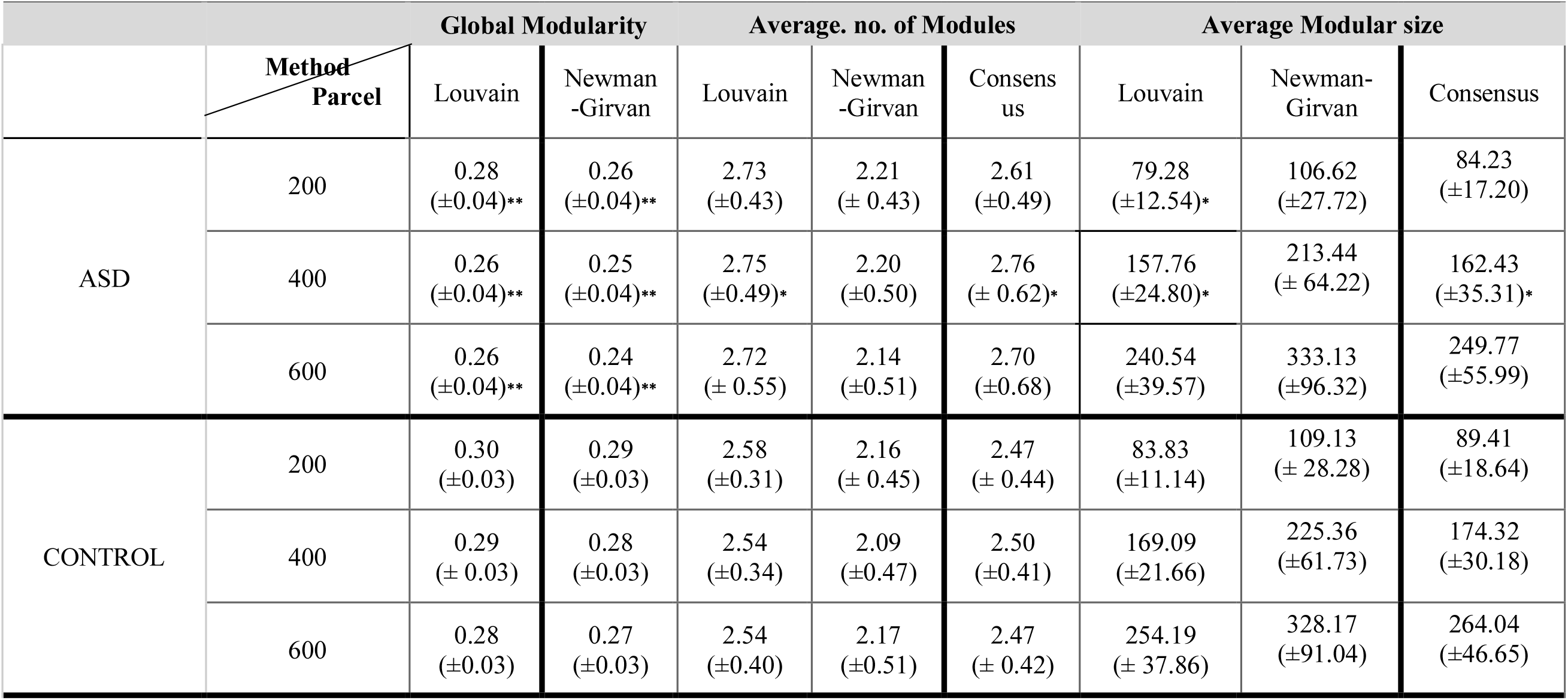
Group difference in global modularity measures across atlas parcellations and community detection methods. A significant group difference in global modularity was observed across all the parcellations and community detection methods (*p>0.05, **p>0.001). A significant difference in average modular size and number of modules was observed for Louvain and consensus agreement methods for parcellation size – 400. Non-significant results also showed similar group differences across all the parcellations and community detection methods. Statistical test: The group difference for global modularity measures was calculated using permutation test (10,000 iterations, p<0.05)

**Supplementary table 2.**
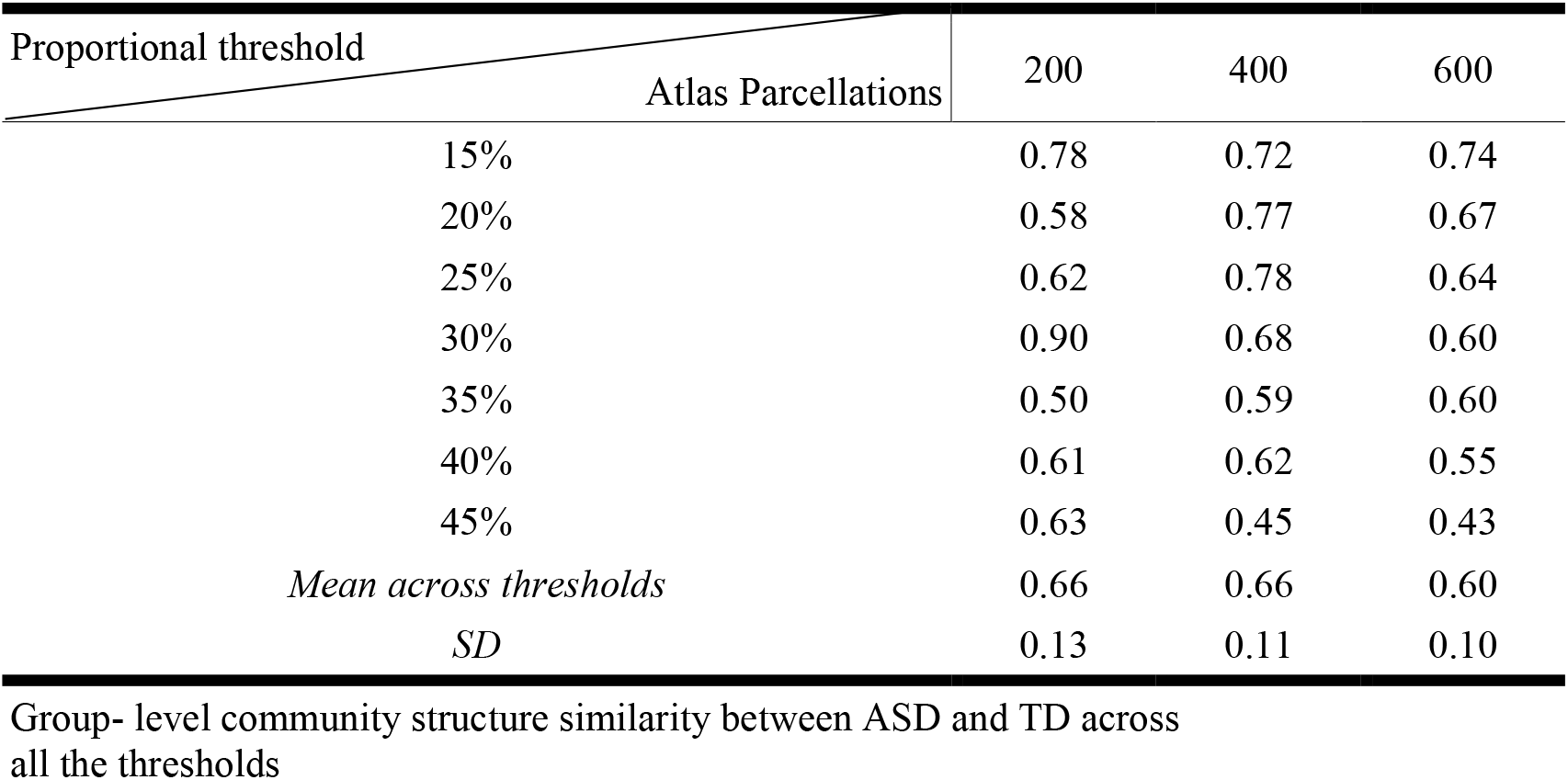
NMI values between ASD and TD across atlas parcellations.

**Supplementary table 3.**
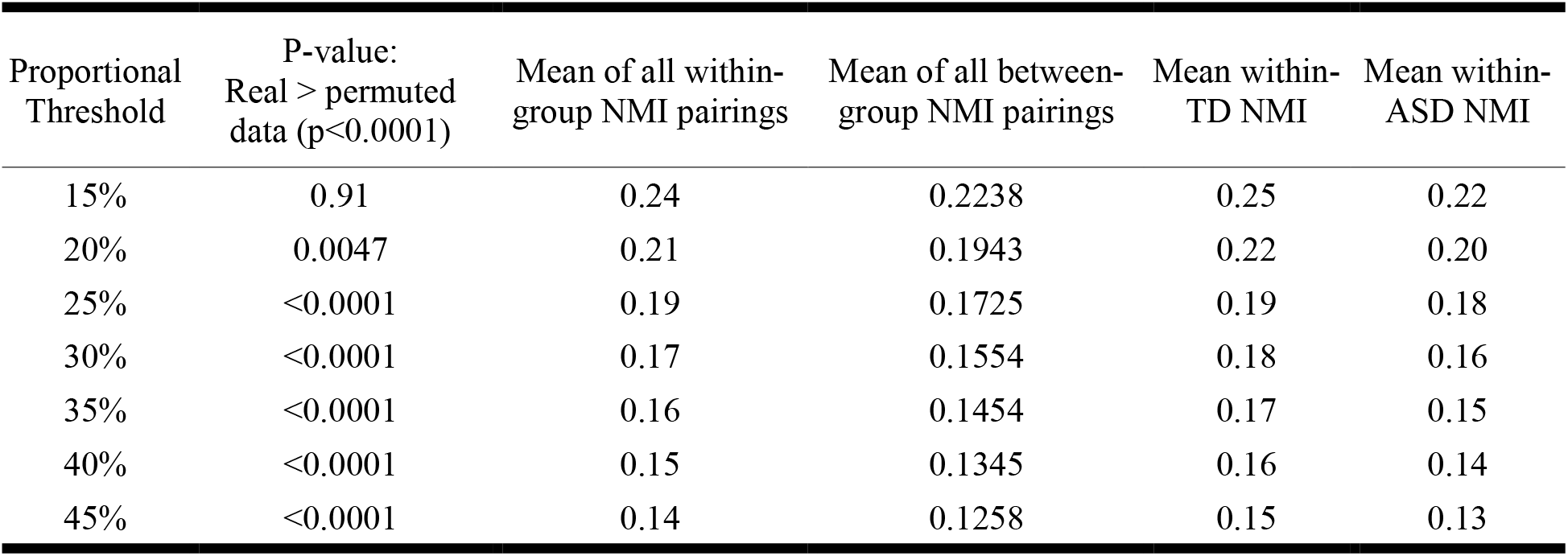
NMI permutation testing between ASD and TD group (Atlas parcellation 400)

**Supplementary table 4.** NMI permutation testing between ASD and TD group (Atlas parcellation 600)

| Proportional Threshold | P-value:<br>Real > permuted<br>data (p<0.0001) | Mean of all within-group NMI pairings | Mean of all between-group NMI pairings | Mean within-TD NMI | Mean within-ASD NMI |
| --- | --- | --- | --- | --- | --- |
| 15% | 0.91 | 0.21 | 0.20 | 0.23 | 0.20 |
| 20% | 0.0014 | 0.18 | 0.17 | 0.20 | 0.17 |
| 25% | <0.0001 | 0.16 | 0.15 | 0.18 | 0.15 |
| 30% | <0.0001 | 0.15 | 0.14 | 0.16 | 0.14 |
| 35% | <0.0001 | 0.14 | 0.13 | 0.15 | 0.13 |
| 40% | <0.0001 | 0.13 | 0.12 | 0.15 | 0.12 |
| 45% | <0.0001 | 0.13 | 0.11 | 0.14 | 0.11 |

Subject-level within group community structure similarity across all the proportional thresholds Permutation test with 10,000 iterations was performed with FDR correction.

**Supplementary figure 1:**
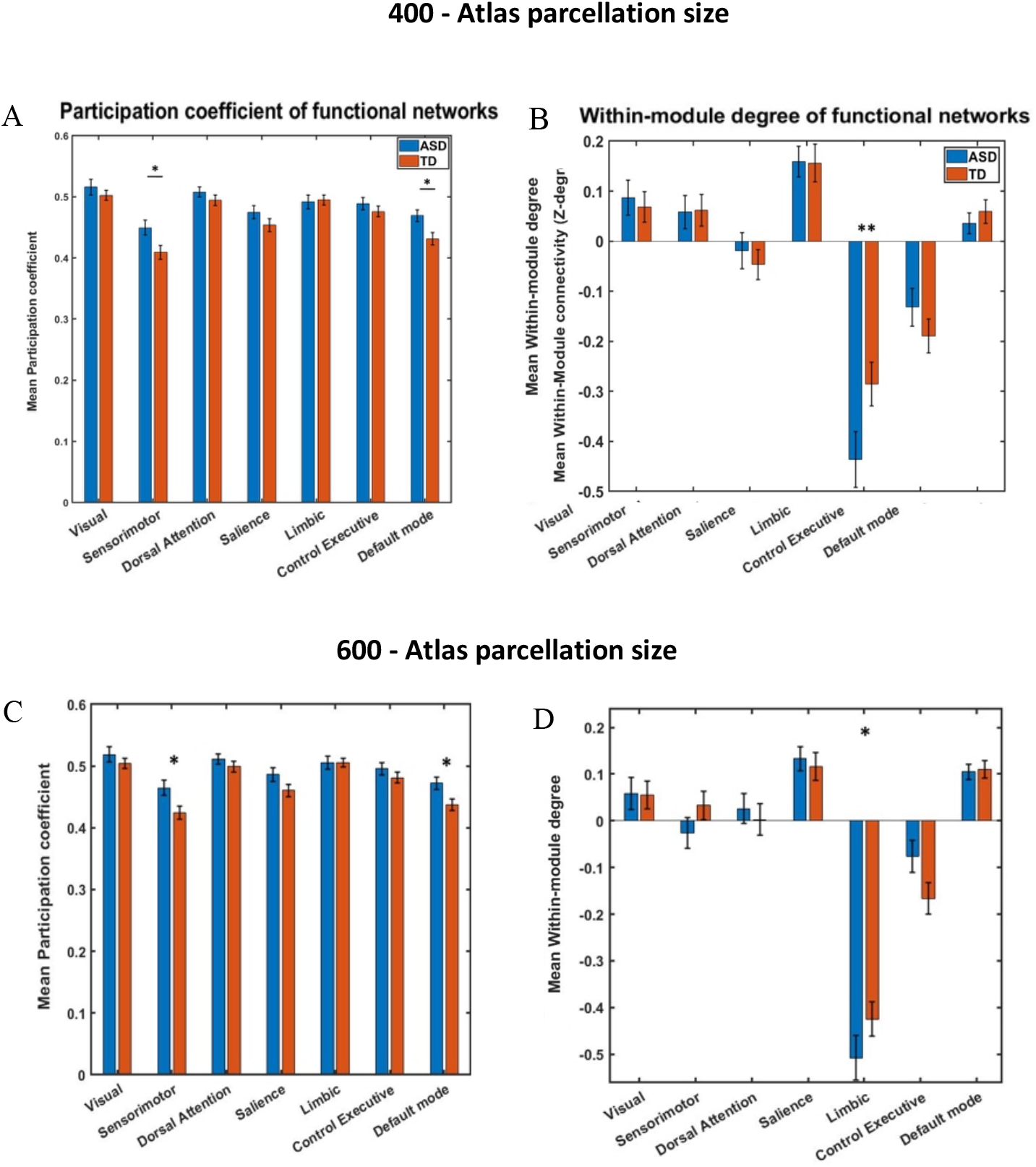
Group difference in modular connectivity across atlas parcellations. A & C. The network-level between–module connectivity was calculated by averaging the PC scores of all the nodes within each network. Sensorimotor and Default-mode network show significantly (FDR corrected, p<0.05) higher between-module connectivity in ASD group as compared to TD group for both the atlas parcellations. B & D. The network-level within-module connectivity revealed that limbic network has significantly (FDR uncorrected, p<0.05) reduced intra- modular connectivity in ASD group for both the atlas parcellations.

**Supplementary figure 2:**
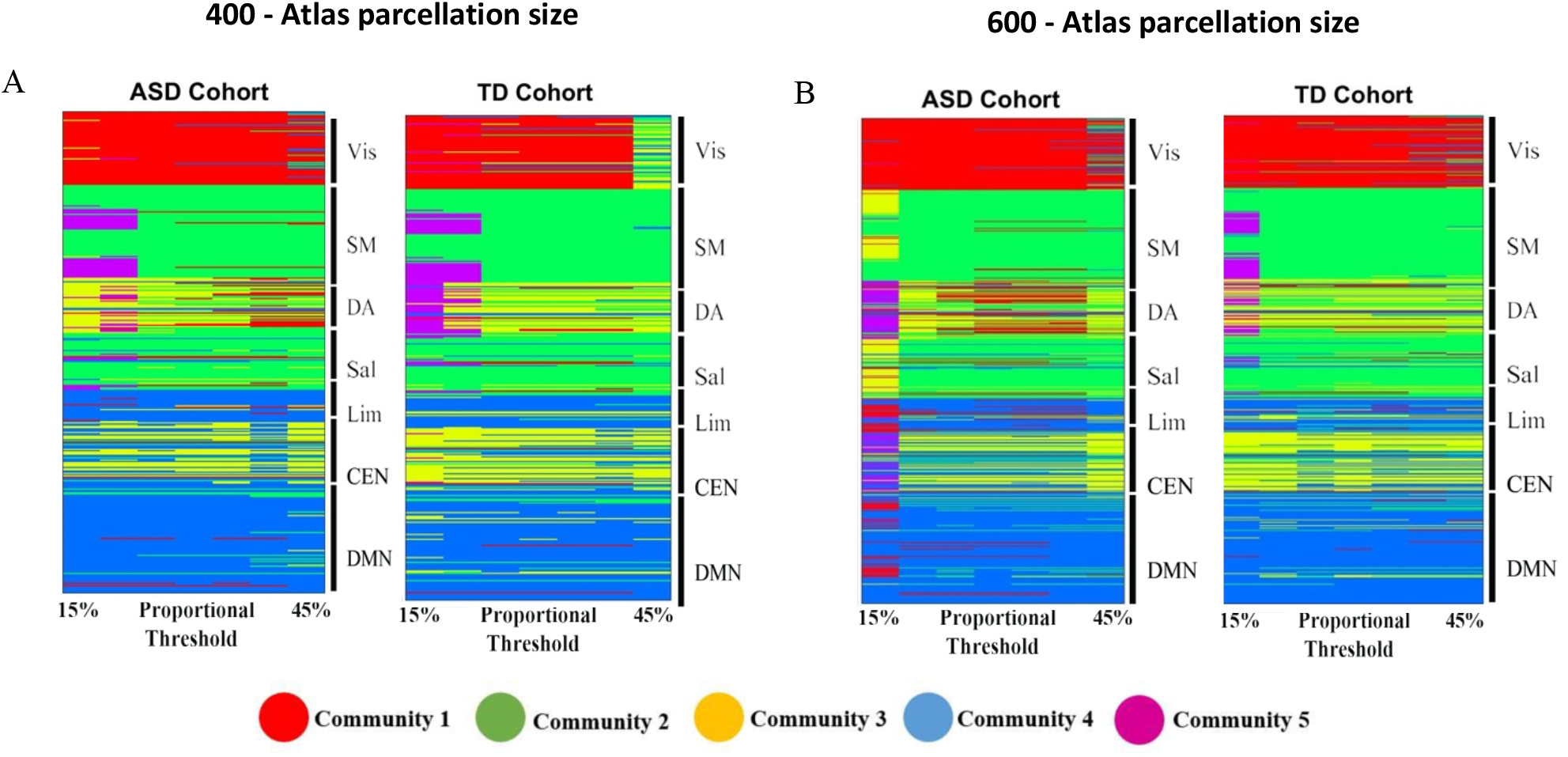
Group-level difference in the community structure revealed overall similarity between ASD and TD groups. The networks are on the vertical axis and graph densities are on the horizontal axis. Regions are coloured by the community assignments of the ROIs. A. No significant group difference in the community structure (supplementary table no. 2) was observed between ASD and TD groups across both the parcellations (A – atlas parcellation size 400; B – atlas parcellation size 600)

**Supplementary figure 3:**
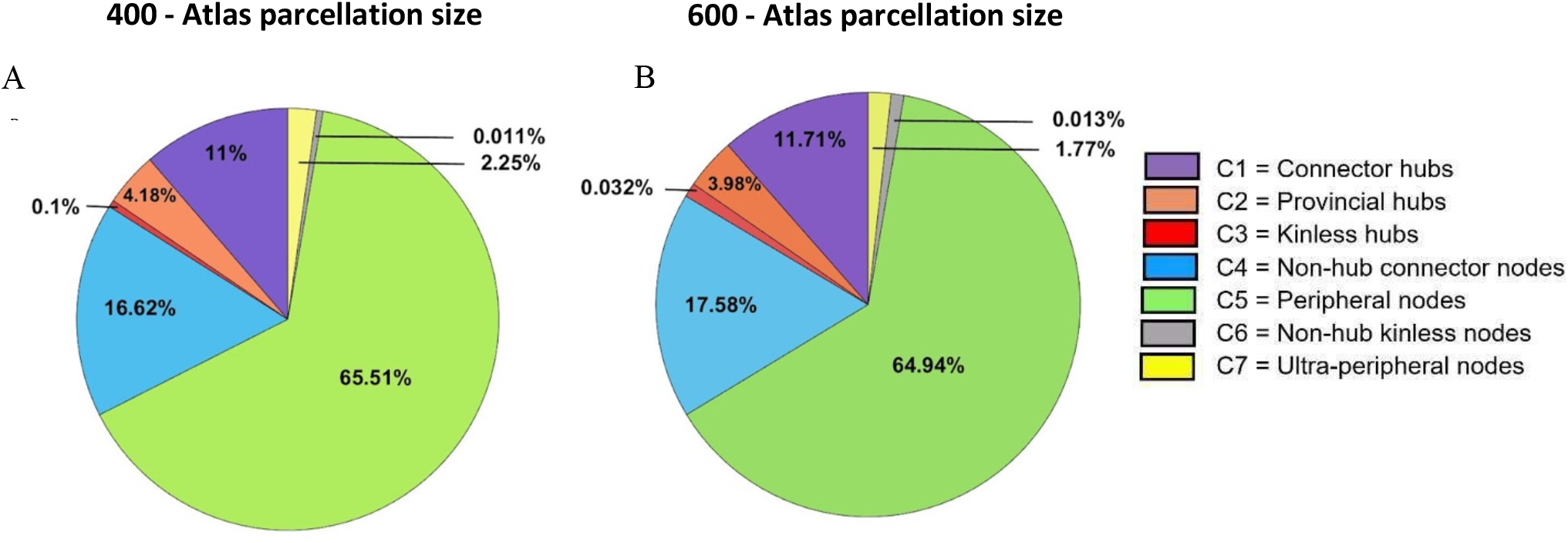
Functional cartography exhibited altered proportion of different node types in ASD group. A & B Whole-brain group-averaged proportions of node types in ASD group shown for two atlas parcellations– 400 & 600 respectively. This was calculated separately on binarized and proportionally thresholded graphs (15%, 20%, 25%, 30%, 35%, 40%, and 45%). For each subject, individual node-type proportion was calculated by averaging across the six thresholds

## Notes

### Competing Interest Statement

The authors have declared no competing interest.

### Summary of Updates

The abstract, part of the introduction and discussion section is modified in the revised version. The supplementary information is also updated.

